# Integrating dedicated and opportunistic surveys to estimate seasonal common dolphin density off mainland Portugal

**DOI:** 10.64898/2026.09.23.753696

**Authors:** Moritz Klaassen, Finn Lindgren, Marc Fernandez, Andrew Houldcroft, Miguel P. Martins, Ana M. Correia, Iolanda M. Silva, Nuno Oliveira, Andreia Torres-Pereira, Ana Marçalo, Sara Martino, Tiago A. Marques, Filipe Alves

## Abstract

1. Common dolphins (*Delphinus delphis*) are the most abundant cetaceans off mainland Portugal and a species of conservation and bycatch concern, so reliable abundance estimates are a management priority. Yet no single survey provides adequate spatiotemporal coverage: dedicated programmes offer sparse snapshots, while opportunistic programmes are spatially constrained by their routes. Integrating them is methodologically demanding, and the existing methods that do so in a single model estimate the density of animal groups, not of the individual animals that management often requires.
2. We develop a spatiotemporal marked log-Gaussian Cox process that integrates four structurally distinct surveys through a shared intensity surface, comprising environmental covariates, a latent spatial field, and survey-specific detection functions. Group size enters as a zero-truncated negative binomial mark with its own spatiotemporal field. We apply it to a dedicated aerial survey and three opportunistic ship-based programmes off mainland Portugal, comprising 162,514 km of on-effort transects and 1,756 sightings of 17,000 animals. We fit the model in a Bayesian framework with integrated nested Laplace approximations, yielding seasonally resolved group and animal density surfaces across the Portuguese Exclusive Economic Zone, and abundance estimates with fully propagated uncertainty.
3. Predicted animal density concentrated over the shelf and upper slope in every season, peaking northwest, and increased in shallower, cooler, more productive waters and over steeper seabed. Abundance nearly doubled from summer (77,000 [95% credible interval: 60,000, 98,000]) to winter (146,000 [99,000, 208,000]). Seasons shared 56% of the spatial field’s variance, so effort from any programme in any season informed all others, and the seasonal contrast remained estimable despite uneven coverage.
4. *Synthesis and applications.* Our framework allows managers to combine dedicated surveys with existing platforms of opportunity to estimate animal density by season. Because group size is modelled explicitly, densities are estimated in animals rather than groups, as required for bycatch risk assessment and marine spatial planning. Because each survey enters through its own observation model, the approach accommodates programmes that differ in platform and protocol and transfers to any species or region where several surveys each cover part of a population.

## 1 Introduction

Accurate estimates of wildlife abundance and distribution are required for a wide range of management actions. For the common dolphin (*Delphinus delphis*) in the Northeast Atlantic, this need is particularly acute for managing bycatch in fishing gear, the most critical anthropogenic threat to the species (Murphy et al., 2021). Bycatch risk depends on where and when fishing effort and animals overlap, and neither is fixed: common dolphin distribution in Iberian and Biscay waters shifts within and between seasons (Lambert et al., 2022), and bycatch mortality in the Bay of Biscay is concentrated in winter (ICES, 2023a). Off mainland Portugal, where seasonal upwelling sustains dense stocks of small pelagic fish (Santos et al., 2001), the common dolphin is the most abundant cetacean species (Gilles et al., 2023). Its diet is dominated by sardines (*Sardina pilchardus*) (Marçalo et al., 2018; Silva, 1999), the same stock that supports the country’s largest coastal fishery. Dolphins and fishing gear come into contact across several fisheries (DGRM, 2026b). Fixed nets, including gillnets and trammel nets, are the most consistent source of cetacean bycatch nationwide and fish year-round (Alexandre et al., 2022; Marçalo et al., 2024; Vingada & Eira, 2018). Purse-seine interactions are tied to the sardine fishery (Marçalo et al., 2015), in which the common dolphin accounted for 89% of cetacean–fishery interaction events over a 15-year observer record and was the only species with documented mortality (Dias et al., 2022).

The management response is organised around when and where this mortality occurs. The International Council for the Exploration of the Sea (ICES) advises keeping bycatch mortality below a removal limit derived with the potential biological removal approach, the maximum number of animals that can be removed each year without depleting the population, and recommends seasonal fishery closures (ICES, 2023a). Winter closures of this kind have been in force in the neighbouring Bay of Biscay since 2024 and are prescribed in EU law through 2026 (European Commission, 2025). Trends in the abundance and distribution of the species additionally feed into assessments of Good Environmental Status under the EU Marine Strategy Framework Directive (Murphy et al., 2021). Removal limits are counted at the individual-animal level, closures must be placed where and when animals concentrate, and status assessments track abundance through time. Therefore, all require absolute animal density, resolved by season, rather than a relative index or a single season estimate.

For common dolphins off mainland Portugal, however, no survey programme has to date delivered absolute animal density resolved by season. The surveys able to deliver absolute abundance follow randomised designs analysed with distance sampling (Buckland et al., 2001); off Atlantic Europe, these are the SCANS (Small Cetaceans in European Atlantic waters and the North Sea) aerial and shipboard surveys, conducted four times since 1994, always in summer (Gilles et al., 2023; Hammond et al., 2002, 2013). Each provides a detailed spatial snapshot, but cannot resolve the seasonal redistribution that the management measures above depend on; the resulting lack of seasonal information at assessment-unit scale is identified as a knowledge gap in the OSPAR regional status assessment (Geelhoed et al., 2022). Monitoring programmes that run through the rest of the year exist, but were built for a different purpose and reach only part of the domain: platforms of opportunity follow fixed commercial routes rather than a randomised design, with documented biases (Oliveira-Rodrigues et al., 2022), and ship effort is often concentrated on the shelf (Martins et al., 2026). This is not a peculiarity of common dolphins in Portuguese waters: line-transect coverage of cetaceans is sparse and uneven in space and time across most of the world’s oceans (Kaschner et al., 2012), and most populations are monitored through a small number of partial programmes.

Statistical data integration offers a way around this trade-off. The underlying true animal distribution can be modelled as a single latent surface observed differently by each programme, letting effort from every programme in every season inform the whole spatiotemporal domain (Isaac et al., 2020; Miller et al., 2019). Multi-survey integration of distance-sampling data has so far been developed mainly within the classical two-stage approach (Hedley & Buckland, 2004), in which detection functions are fitted first, and spatial density is modelled second (Miller et al., 2021). Point processes offer a one-stage alternative, in which sightings are modelled as a log-Gaussian Cox process (LGCP) thinned by imperfect detection and the detection function and intensity surface are estimated jointly (Møller et al., 1998; Yuan et al., 2017). In this joint hierarchical model, detection uncertainty propagates directly into density and abundance, and a randomised survey can set the absolute scale of the predictions while non-randomised programmes can inform the spatial and seasonal patterns.

Within this point-process formulation, two components that management-ready density estimation requires have not yet been brought together. First, detection conditions vary within as well as between programmes, and detection covariates, standard practice in classical distance sampling (Marques & Buckland, 2003), have to our knowledge not been carried into the point-process formulation across multiple integrated surveys. Second, and more fundamentally, points of this process are sightings, and for a group-living species such as the common dolphin, a sighting is a group of animals, not an individual. The group-to-animal step remains underexplored: group size was explicitly left as a future extension in the original point-process formulation of distance sampling (Yuan et al., 2017), and has since been modelled jointly with the intensity surface for a single terrestrial survey (Houldcroft et al., 2025), but not within an integrated multi-survey model or at seasonal resolution. This gap matters because a group surface and an animal surface do not need to agree: where group size varies in space, and in time over space, as it does for many social species, a model of groups alone can misstate where animals concentrate.

Here, we develop a spatiotemporal marked LGCP that closes both gaps: all surveys observe one shared intensity surface, each through its own detection function, and group size enters as a spatiotemporally varying mark that converts group density to animal density. We apply it to four monitoring programmes off mainland Portugal, comprising a dedicated aerial survey and three ship-based programmes. We use the fitted model to (i) estimate seasonal group and animal density and abundance for the Portuguese Exclusive Economic Zone (EEZ), (ii) identify the environmental conditions associated with high dolphin density, and (iii) characterise the seasonal spatial patterns of animal density, the quantity required to evaluate seasonal closures, assess bycatch risk and inform marine spatial planning (Hazen et al., 2018; Maxwell et al., 2015).

## 2 Materials and Methods

### 2.1 Study area and survey programmes

The study area is the EEZ of mainland Portugal, on the western Iberian shelf and slope within the Canary/Iberian Eastern Boundary Upwelling System. We integrated four line-transect programmes covering this area (Table 1). They differ in platform, search protocol, truncation width and the way perpendicular distance was obtained, and each therefore enters the model through its own detection function.

**Table 1:** Survey programmes integrated in the analysis, with years and seasons of coverage, truncation half-width (*W*), on-effort transect length, and common dolphin (*Delphinus delphis*) sightings entering the model.

| Programme | Platform | Years | Seasons | $W$ (km) | Effort (km) | Groups | Animals |
| --- | --- | --- | --- | --- | --- | --- | --- |
| SPEA | Ship | 2004–2026 | All | 0.3 | 100,607 | 1,177 | 10,216 |
| CETUS | Ship | 2012–2024 | Spring–Autumn | 1.0 | 52,564 | 334 | 4,354 |
| ATLANTIDA | Ship | 2021–2024 | All | 0.5 | 1,845 | 67 | 540 |
| SCANS-IV | Aircraft | 2022 | Summer | 0.4 | 7,498 | 178 | 1,890 |
| All programmes |  | 2004–2026 | All |  | 162,514 | 1,756 | 17,000 |

The longest of the four is run by Sociedade Portuguesa para o Estudo das Aves (SPEA) which follows the European Seabirds at Sea methodology (Tasker et al., 1984). Seabirds are the primary target and cetaceans are recorded alongside. Vessels were research ships of the Portuguese Institute for Sea and Atmosphere used during pelagic fish stock assessments, together with smaller motor and sailing vessels and campaigns. Experienced observers searched on one side of the vessel and recorded perpendicular distance in four bands out to 300 m using per-observer range sticks calibrated in the field.

Two further programmes are run from Portuguese vessels under different protocols. CETUS has monitored cetaceans from cargo and oceanographic vessels used as platforms of opportunity since 2012 (Oliveira-Rodrigues et al., 2022), while ATLANTIDA surveys the northern coast of Portugal from a 12 m catamaran with an elevated observation deck. Both programmes search both sides of the vessel ahead of the route with at least two observers. CETUS additionally scores each observer’s experience, which we used as a detection covariate because observer experience is the observer-level bias documented for this programme (Oliveira-Rodrigues et al., 2022).

The fourth is the Portuguese component of the pan-European SCANS-IV aerial survey, flown in July and August 2022 (Gilles et al., 2023). It is the only one of the four designed specifically to estimate cetacean abundance, and the only one with a randomised survey design: transects were placed within blocks to give equal coverage probability, using equal-spaced zigzag designs offshore and parallel designs in coastal blocks, so that every point within a block has the same probability of being surveyed. We excluded records from circle-back procedures, which resurvey a segment of transect after a detection to estimate detection on the trackline.

Coverage across programmes is uneven in both space and time, and no single programme spans the domain in all seasons (Fig. 1). SPEA contributes the longest time series, covers all four seasons and provides nearly all of the winter effort, but is concentrated on the shelf, with offshore waters surveyed at much lower intensity. CETUS reaches comparably far offshore, though only along fixed routes of its host vessels. ATLANTIDA covers only a short stretch of the northern coast, in all seasons but at low effort. SCANS-IV is the only programme to cover the offshore domain with spatially balanced effort, but for a single summer. Integrating them is therefore what allows density to be resolved seasonally across the full space–time domain rather than in the subset any one programme reaches.

**Figure 1.**
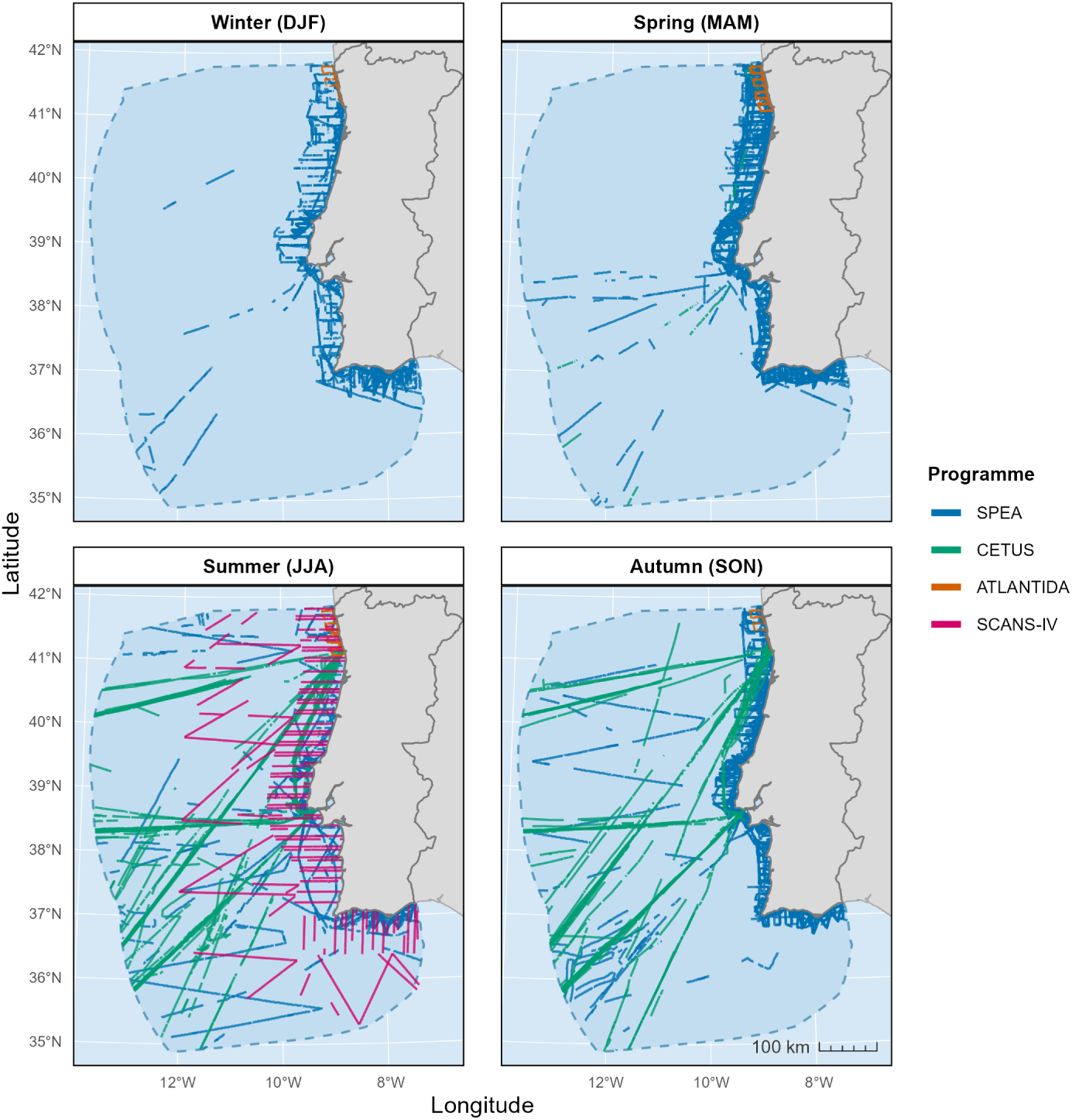
Spatial and seasonal distribution of survey transects within the mainland Portuguese Exclusive Economic Zone (EEZ), by programme: SPEA, CETUS, ATLANTIDA, and SCANS-IV. Effort is shown separately for winter (DJF), spring (MAM), summer (JJA), and autumn (SON).

### 2.2 Environmental covariates

Candidate covariates were derived from CMEMS IBI reanalysis products (sea surface temperature, salinity, mixed-layer depth, bathymetry) (CMEMS, 2020b), CMEMS ocean colour (chlorophyll-*a*) (CMEMS, 2022), CMEMS IBI biogeochemistry (zooplankton biomass) (CMEMS, 2020a), Natural Earth coastline (Natural Earth, 2025), and EMODnet geomorphology (canyon and seamount polygons) (EMODnet Geology, 2022). All layers were resampled to a common 0.027° grid matching the coarsest native spatial resolution among the raster datasets. Seasonal variables were aggregated to seasonal climatologies (winter, spring, summer, autumn) over 2004–2026, matching the span of the survey effort. Slope and an analogous sea-surface-temperature gradient were derived from bathymetry and SST via terrain analysis; distance to coast, canyon, and seamount were computed as Euclidean distances in an equal-area projection. Of eleven candidate covariates, salinity, mixed-layer depth, and zooplankton were dropped for collinearity (correlation and variance inflation factor (VIF)), leaving eight from which the final set was chosen by backward elimination (§2.4). All covariates were standardised before fitting.

### 2.3 Statistical model

#### 2.3.1 Model formulation

For each survey programme *k*, we model the occurrence of dolphin groups as a log-Gaussian Cox process with intensity

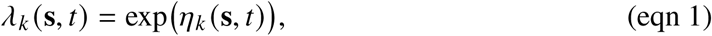

where **s** ∈ D ⊂ R^2^ is a spatial location, *t* ∈ {1, . . . , 4} is a season (winter, spring, summer, autumn), and *η_k_* (**s**, *t*) is a programme-specific linear predictor sharing all structure across programmes except a programme offset,

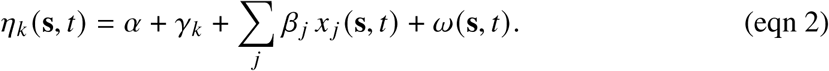

Here *α* is a shared intercept, *γ_k_* is a sum-to-zero IID offset across the four programmes absorbing residual between-programme differences, *β_j_* are linear coefficients for standardised environmental covariates *x _j_* (**s**, *t*), and *ω*(**s**, *t*) is a latent Gaussian spatial field replicated across seasons with an exchangeable correlation structure,

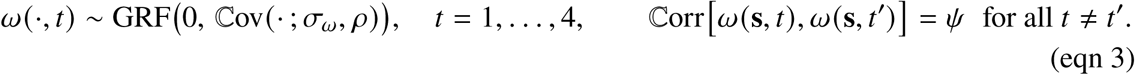

*ω*(**s**, *t*) is a Gaussian random field over continuous space with Matérn covariance, represented through its stochastic partial differential equation (SPDE) approximation on a triangulated mesh (Lindgren et al., 2011), which yields a sparse Gaussian Markov random field (Appendix S1.3). *σ_ω_* is the marginal standard deviation of the field and *ρ* its practical range. Priors on (*ρ*, *σ_ω_*) follow the penalised-complexity (PC) approach (Simpson et al., 2017), with P(*ρ* < 50 km) = 0.01 and P(*σ_ω_* > 1) = 0.05. The exchangeable structure imposes a correlation *ψ* between every pair of seasons, which allows seasons with sparse effort to borrow strength from those with more.

#### 2.3.2 Detection functions

Each programme *k* observes the shared intensity through its own detection function. Let *y*(**s**) denote the perpendicular distance from **s** to the nearest transect line in the strip *C_k_* ⊂ D covered by programme *k*, bounded by truncation half-width *W_k_*. Two detection function forms are used across programmes: the half-normal,

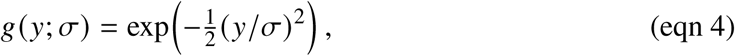

and the hazard-rate,

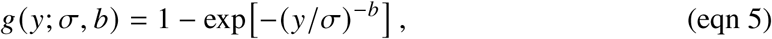

with *σ* > 0 the scale parameter and, for the hazard-rate model (Eq. 5), *b* > 0 the shape parameter. Where the detection scale depends on covariates, it enters on the log scale as a linear predictor, following the multiple covariate distance sampling formulation of Marques and Buckland (2003).

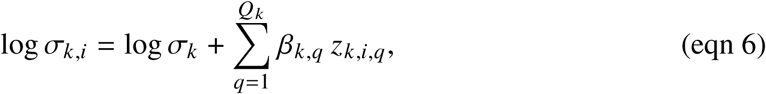

where *σ_k_* is the scale parameter at the reference value of the covariates for programme *k*, *z_k,i,q_* is the value of the *q*-th detection covariate for programme *k* associated with detection *i*, and *β_k_*_,*q*_ is its programme-specific coefficient. The covariate candidate sets differ by programme, reflecting what each records on effort. Continuous covariates were standardised and entered linearly on the log scale; categorical covariates entered as factor contrasts. For each programme we fitted both key functions with candidate covariates in turn, and both with no covariate, and compared the resulting models by Deviance Information Criterion (DIC). DIC guided the comparison but did not settle it. For SPEA, we retained the half-normal on the plausibility of the fitted shape rather than the lower-DIC hazard-rate, following the selection made for the same protocol by Martins et al. (2026). Full comparisons are given in Appendix S1.

#### 2.3.3 Point-process likelihood

The log-likelihood contribution of programme *k* is

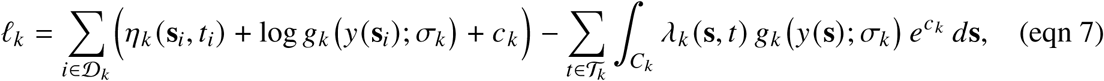

where *D_k_* indexes the detections of programme *k*, *T_k_* ⊆ {1, . . . , 4} is the set of seasons in which programme *k* was active, *C_k_* denotes the one-sided (folded-distance) covered strip [0, *W_k_*] over which programme *k*’s effort is defined, and *c_k_* = log 2 accounts for two-sided transect protocols (all programmes except SPEA, which searches one side only).

#### 2.3.4 Group size

Each sighted group carries a mark *G_i_* ≥ 1, the observed group size. Because groups are only ever recorded conditional on being sighted, we model the observed marks as draws from a zero-truncated negative binomial,

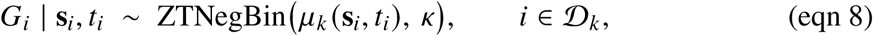

where *μ_k_* (**s**, *t*) is the mean of the untruncated negative binomial and *κ* is a shared overdispersion parameter. The untruncated mean is modelled on the log scale as

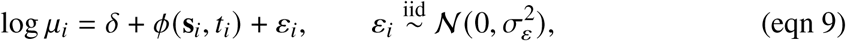

with its own intercept *δ*, its own latent spatial field *φ*(**s**, *t*), and a sighting-level random effect *ε_i_*. The field *φ* uses the same Matérn SPDE construction as *ω* (Eq. 3) and is likewise replicated over the four seasons, but its seasonal replicates are treated as independent. Its PC priors are P(*ρ_φ_* < 30 km) = 0.01 and P(*σ_φ_* > 1) = 0.05. The sighting-level effect *ε_i_* and the negative binomial parameter *κ* describe overdispersion at different levels: *κ* applies to all groups sharing a location and season, whereas *ε_i_* absorbs individual groups that depart from the local mean.

Because *G*≥1 by construction, the observable mean follows by removing the zero mass and renormalising. Writing 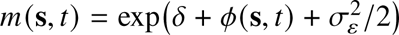 for the mean of *μ_i_* at (**s**, *t*) after marginalising *ε_i_*,

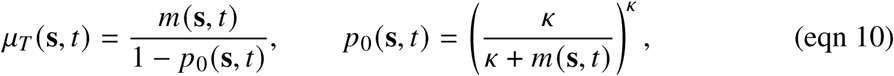

where *p*_0_(**s**, *t*) is the zero mass of the untruncated distribution.

#### 2.3.5 Animal density and abundance

Equation 2 makes the intensity programme-specific, so reporting a single density surface requires choosing a reference programme. We predict at *γ*_SCANS_, the offset of the only randomised survey designed for abundance estimation. Animal density combines the survey-marginal group-level intensity with the truncated mean group size,

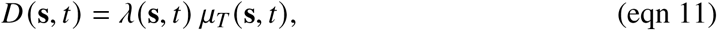

Seasonal abundance follows from integrating animal density over the study domain,

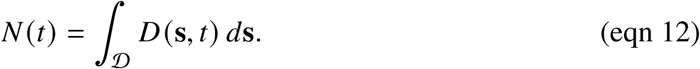

Because groups arise from a conditional Poisson process, the total variance of *N* (*t*), decomposes into a within-realisation term and a field/estimation-uncertainty term (derivation in Appendix S3),

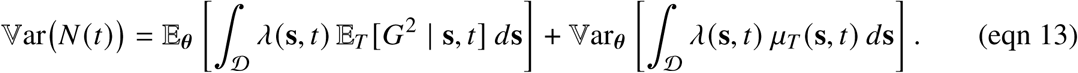

### 2.4 Model selection and assessment

Intensity covariates *x _j_* (**s**, *t*) were selected by backward elimination from the full candidate set of standardised environmental covariates described in §2.2: starting from the model containing all candidates, the covariate whose removal produced the lowest DIC was dropped at each step, and elimination continued until no remaining removal improved DIC.

We compared summer predictions with the published design-based SCANS-III and SCANS-IV block estimates, and seasonal predictions with the model-based estimates of Martins et al. (2026), which were derived from a subset of the SPEA data used here (Appendix S2).

### 2.5 Implementation

All analyses were conducted in R version 4.4.3 (R Core Team, 2025), with models fitted using INLA (version 25.3.24) (Rue et al., 2009) and inlabru (version 2.14.1.9012) (Bachl et al., 2019). All predictions, including fitted detection functions, density surfaces and seasonal group and animal abundances were computed from 2,000 samples of the joint posterior distribution.

## 3 Results

### 3.1 Detection

We retained a half-normal key function for SPEA and hazard-rate key functions for CETUS, ATLANTIDA and SCANS-IV, with sea state on the detection scale for SPEA and SCANS-IV, observer experience for CETUS and no covariate for ATLANTIDA (full model comparisons in Appendix S1). Detection probability fell with increasing sea state on both programmes that carried it and rose with observer experience (Fig. 2).

**Figure 2.**
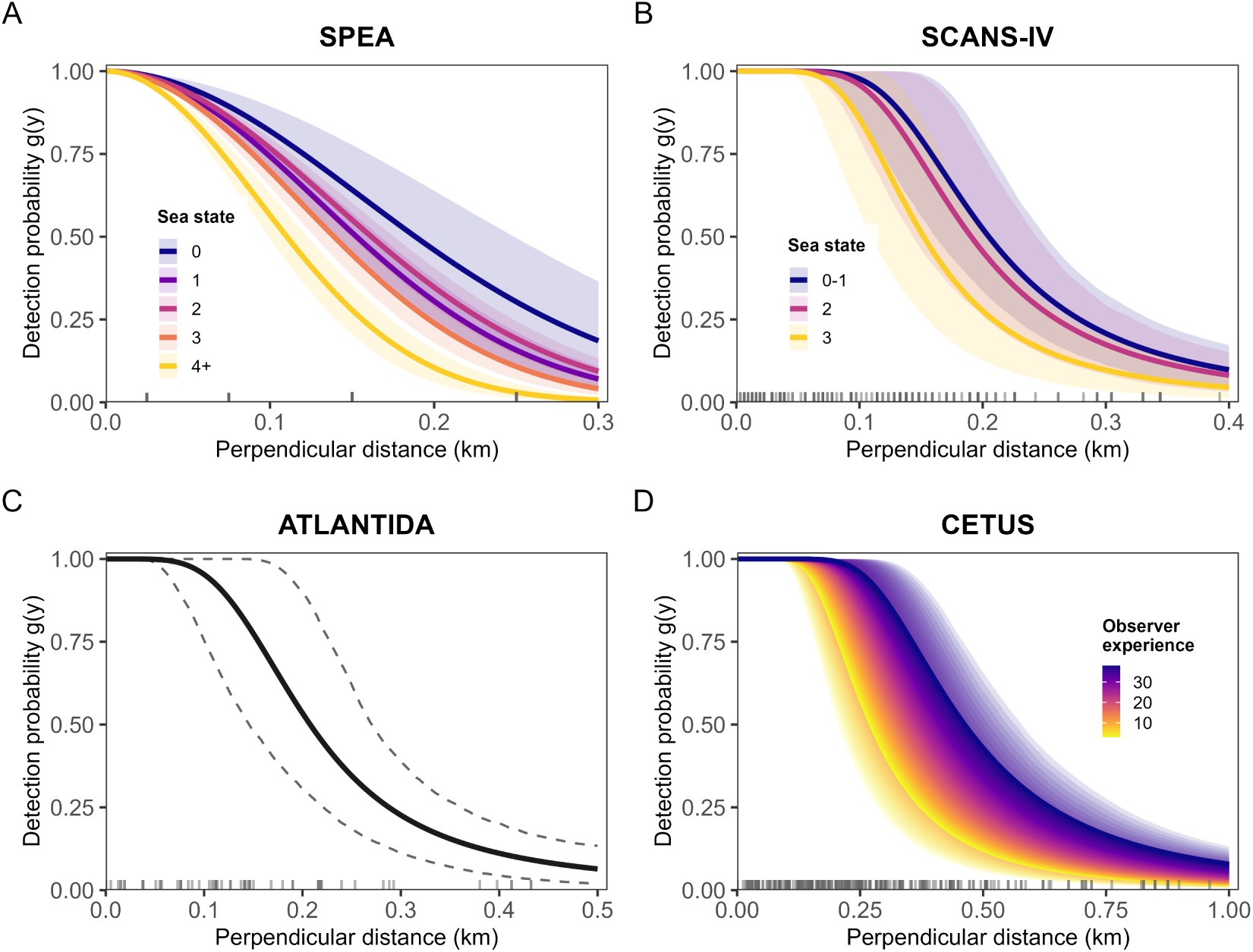
Fitted detection functions by survey programme. (A) SPEA: half-normal detection function by sea state. (B) SCANS-IV: hazard-rate detection function by sea state. (C) ATLANTIDA: hazard-rate detection function. (D) CETUS: hazard-rate detection function by observer experience. Shaded ribbons show 95% credible intervals in all panels; tick marks along each x-axis show the distribution of observed perpendicular distances.

Reference-level effective strip half-widths ranged more than two-fold across programmes, from 0.188 km [95% credible interval: 0.150, 0.221] for SPEA to 0.428 km [0.386, 0.470] for CETUS, with ATLANTIDA at 0.236 km [0.182, 0.300] and SCANS-IV at 0.228 km [0.195, 0.267] (Table 2).

**Table 2:**
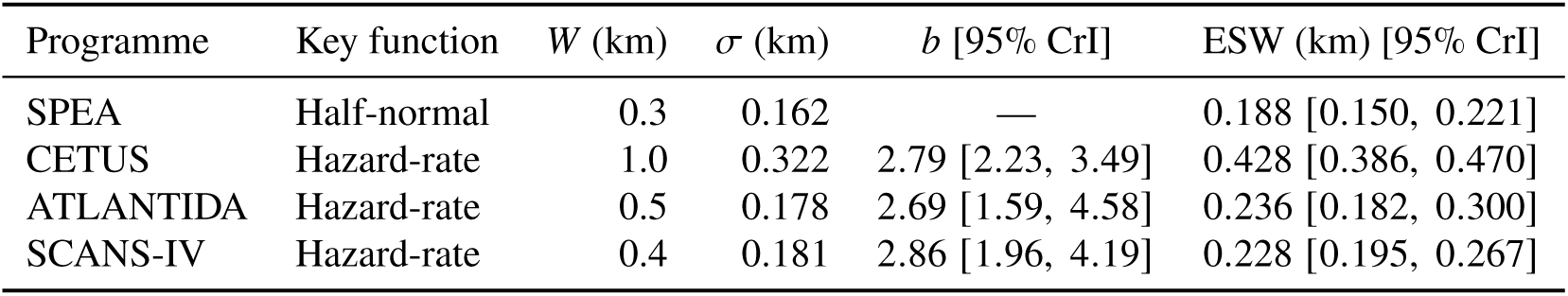
Detection function parameters by survey programme. *W* is the truncation distance, *σ* the scale parameter and *b* the hazard-rate shape parameter, both evaluated at the reference sea state and mean observer experience. ESW is the effective strip half-width. Values in brackets are 95% credible intervals (CrI).

### 3.2 Group density and group size

Four of the five environmental covariates had 95% credible intervals excluding zero (Fig. 3). Group density declined with bathymetry (−0.534 per SD [−0.761, −0.307]) and sea surface temperature (−0.436 [−0.653, −0.219]) and increased with seabed slope (0.287 [0.142, 0.432]) and chlorophyll-*a* (0.099 [0.042, 0.156]); the interval for distance to canyon spanned zero (−0.298 [−0.630, 0.034]).

**Figure 3.**
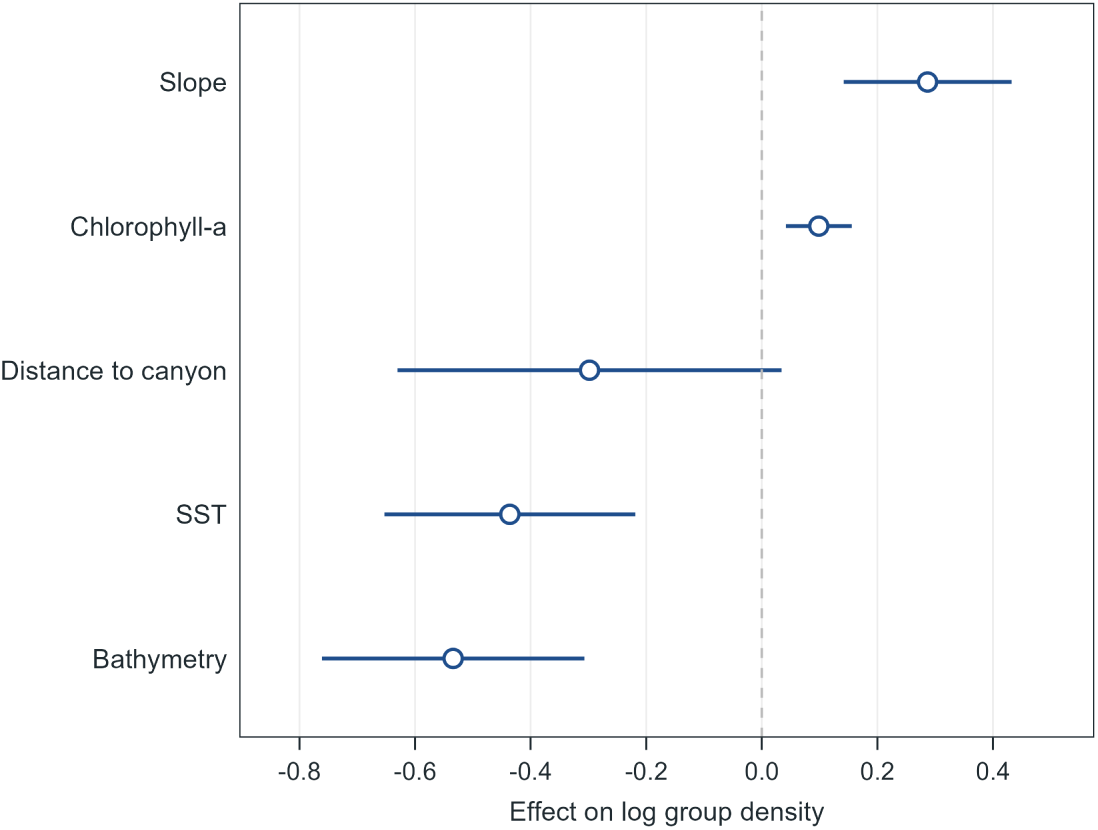
Posterior means and 95% credible intervals for the environmental covariate coefficients. All covariates were standardised, so effects are per standard deviation.

The latent spatial field had a range of 68.7 km [53.6, 87.4] and a marginal standard deviation of 1.073 [0.901, 1.272]; the between-season correlation was *ψ* = 0.558 [0.390, 0.707]. Programme offsets were 0.286 [0.135, 0.438] for SPEA, 0.259 [−0.035, 0.553] for ATLANTIDA, 0.467 [0.265, 0.669] for SCANS-IV and −1.013 [−1.195, −0.830] for CETUS. Fig. 4 decomposes the predicted surface, shown for winter as an example, into the environmental contribution and the latent spatial field, together with the resulting group-and animal-density surfaces: the latent field added coastal structure that the covariates alone did not capture, and scaling by expected group size shifted the main concentration of animals relative to that of groups.

**Figure 4.**
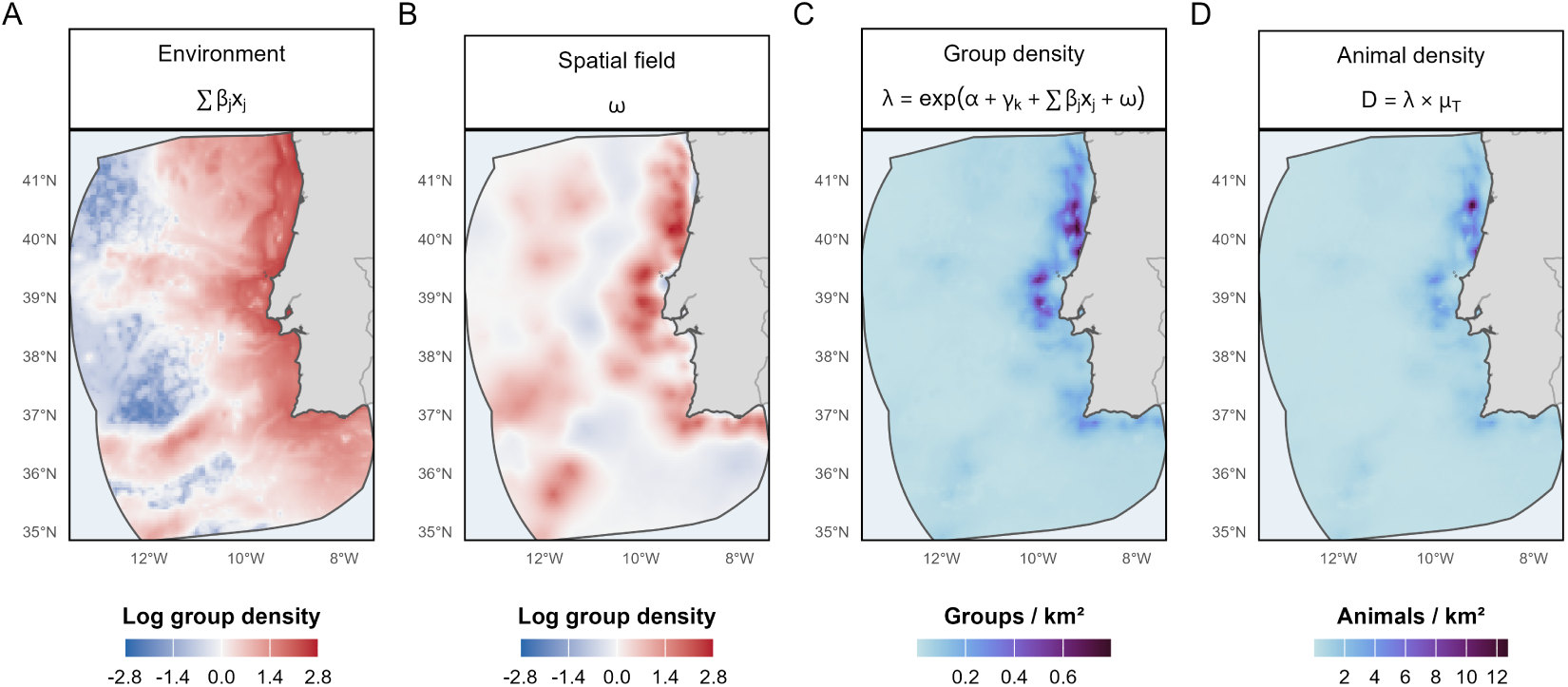
Decomposition of predicted common dolphin (*Delphinus delphis*) density in winter. (A) The environmental contribution to the linear predictor; (B) the latent spatial field; (C) group density; and (D) animal density, obtained by scaling group density by the expected zero-truncated group size. In (C) and (D) the programme offset *γ_k_* is set to the SCANS-IV value.

The group-size intercept was *δ* = 1.936 [1.837, 2.035] on the log scale. Seasonal mean expected group size was stable across the year, ranging from 9.4 to 10.3 animals, while expected group size varied roughly 2.5-fold in space. The group-size field varied at a finer spatial scale than the intensity field, with a range of 37.4 km [21.0, 63.1] against 68.7 km [53.6, 87.4], and had a marginal standard deviation of 0.454 [0.358, 0.563]. The dispersion parameter was *κ* = 2.99 [1.62, 5.44] and the sighting-level standard deviation was *σ_ε_* = 0.714 [0.594, 0.869]. Full posterior summaries are given in Table 3.

**Table 3:** Posterior summaries for the fitted marked log-Gaussian Cox (LGCP) process. Intercepts and coefficients are on the log scale; covariate coefficients *β_j_* are per standard deviation.

| Parameter | Mean | SD | 95% CrI |
| --- | --- | --- | --- |
| **Group intensity** |  |  |  |
| *Fixed effects* |  |  |  |
| Intercept $\alpha$ | -5.213 | 0.193 | [-5.592, -4.834] |
| Bathymetry | -0.534 | 0.116 | [-0.761, -0.307] |
| Sea surface temperature | -0.436 | 0.111 | [-0.653, -0.219] |
| Seabed slope | 0.287 | 0.074 | [ 0.142, 0.432] |
| Chlorophyll-*a* | 0.099 | 0.029 | [ 0.042, 0.156] |
| Distance to canyon | -0.298 | 0.169 | [-0.630, 0.034] |
| *Spatial field ($\omega$)* |  |  |  |
| Range (km) $\rho$ | 68.70 | 8.61 | [53.57, 87.43] |
| Marginal SD $\sigma_\omega$ | 1.073 | 0.094 | [0.901, 1.272] |
| Between-season correlation $\psi$ | 0.558 | 0.081 | [0.390, 0.707] |
| *Programme offsets ($\gamma_k$, sum-to-zero)* |  |  |  |
| SPEA | 0.286 | 0.078 | [ 0.135, 0.438] |
| CETUS | -1.013 | 0.095 | [-1.195, -0.830] |
| ATLANTIDA | 0.259 | 0.151 | [-0.035, 0.553] |
| SCANS-IV | 0.467 | 0.105 | [ 0.265, 0.669] |
| SD of $\gamma_k$ | 0.608 | 0.213 | [ 0.310, 1.136] |
| **Group size** |  |  |  |
| *Fixed effects* |  |  |  |
| Intercept $\delta$ | 1.936 | 0.050 | [1.837, 2.035] |
| *Spatial field ($\phi$)* |  |  |  |
| Range (km) $\rho_\phi$ | 37.36 | 10.81 | [20.96, 63.14] |
| Marginal SD $\sigma_\phi$ | 0.454 | 0.052 | [0.358, 0.563] |
| *Overdispersion* |  |  |  |
| Dispersion $\kappa$ | 2.993 | 0.985 | [1.615, 5.435] |
| Sighting-level SD $\sigma_\varepsilon$ | 0.714 | 0.070 | [0.594, 0.869] |

### 3.3 Abundance and animal density

Estimated abundance in the mainland Portuguese EEZ was highest in winter and lowest in summer, with spring above autumn in between (Table 4). The number of individuals nearly doubled from summer (76,908 [59,505, 97,901]) to winter (146,071 [99,105, 208,218]); spring and autumn estimates were 110,836 [72,896, 163,127] and 86,435 [64,301, 114,451] individuals. Mean densities were 0.465, 0.353, 0.245 and 0.275 individuals per km^2^ in winter, spring, summer and autumn.

**Table 4:** Estimated seasonal abundance of common dolphin (*Delphinus delphis*) groups and individuals within the mainland Portuguese Exclusive Economic Zone (EEZ), and within its continental shelf (waters shallower than 200 m). Values are posterior means with 95% credible intervals (CrI) and coefficients of variation (CV). Density is individuals per km^2^.

| Season | Groups |  | Individuals |  | Density |
| --- | --- | --- | --- | --- | --- |
| | $\hat{N}$ (95% CrI) | CV | $\hat{N}$ (95% CrI) | CV | |
| *Full EEZ* |  |  |  |  |  |
| Winter (DJF) | 15,280 (10,725–21,387) | 0.18 | 146,071 (99,105–208,218) | 0.19 | 0.465 |
| Spring (MAM) | 11,747 (7,995–16,949) | 0.20 | 110,836 (72,896–163,127) | 0.22 | 0.353 |
| Summer (JJA) | 7,649 (6,034–9,497) | 0.12 | 76,908 (59,505–97,901) | 0.13 | 0.245 |
| Autumn (SON) | 8,396 (6,335–10,952) | 0.14 | 86,435 (64,301–114,451) | 0.15 | 0.275 |
| *Shelf (<200 m)* |  |  |  |  |  |
| Winter (DJF) | 4,235 (3,063–5,785) | 0.15 | 40,575 (28,546–56,662) | 0.18 | 1.931 |
| Spring (MAM) | 1,951 (1,471–2,552) | 0.14 | 15,667 (11,526–20,689) | 0.15 | 0.746 |
| Summer (JJA) | 2,408 (1,918–3,020) | 0.12 | 22,826 (17,575–29,542) | 0.13 | 1.087 |
| Autumn (SON) | 3,084 (2,351–4,000) | 0.14 | 30,342 (22,515–39,817) | 0.14 | 1.444 |

On the shelf, waters shallower than 200 m covering 21,012 km^2^ (6.7% of the study area), winter abundance was again nearly twice that of summer, 40,575 individuals [28,546, 56,662] against 22,826 [17,575, 29,542], with the shelf minimum in spring at 15,667 [11,526, 20,689] and autumn at 30,342 [22,515, 39,817] (Table 4). Shelf densities of 1.93 (winter) and 1.09 (summer) individuals per km^2^ were roughly four times the corresponding EEZ-wide values, and the shelf held about 28% of the winter EEZ total.

Predicted density concentrated in a nearshore strip along the west coast in every season, with the offshore domain uniformly low; the winter surface was the most intense, with a coastal hotspot near 40.5^◦^N (Fig. 5A). Fig. 5B illustrates the fine spatial scale at which the fitted model can predict, shown for the northern coast in autumn as an example.

**Figure 5.**
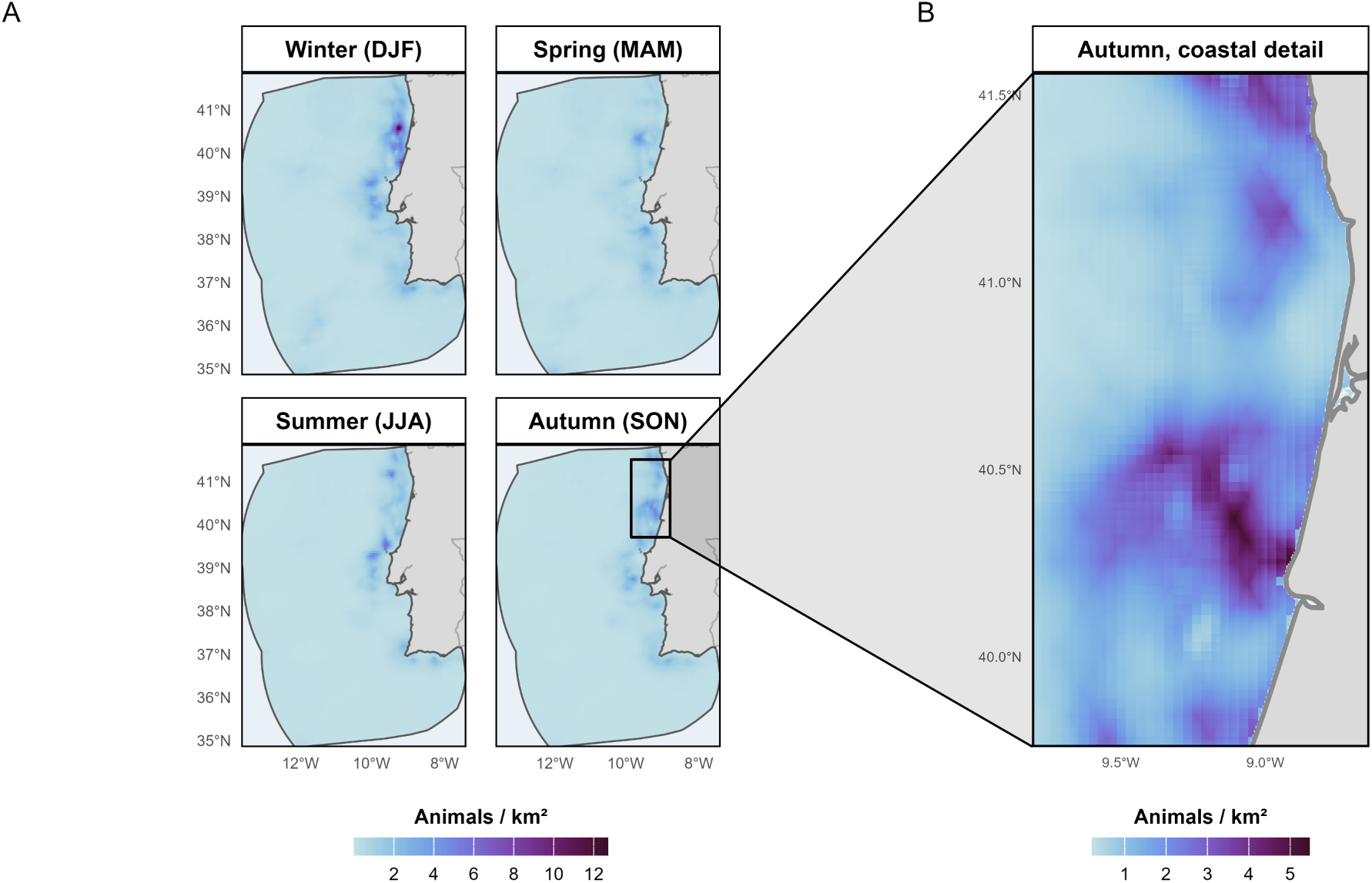
Predicted common dolphin (*Delphinus delphis*) density within the mainland Portuguese Exclusive Economic Zone. (A) Animal density by season. (B) Example of fine-scale density predictions off the northern Portuguese coast in autumn.

## 4 Discussion

Integrating four survey programmes, each through its own observation model, gave seasonally resolved density surfaces for common dolphins across the whole mainland Portuguese EEZ with propagated uncertainty, which no single programme could provide. Abundance was highest in winter and lowest in summer, nearly doubling between the two, and winter remained close to twice summer when the estimate was restricted to the shelf, where the net fisheries operate. Density concentrated on the shelf and upper slope in every season, with the winter maximum on the northern shelf, and increased in shallower, cooler and more productive water and over steeper seabed.

### 4.1 Implications for bycatch management and status assessment

In the Bay of Biscay, where bycatch mortality of common dolphins is concentrated in winter, the mitigation measure now in force is a winter fishery closure, following ICES advice (European Commission, 2025; ICES, 2023a). Off mainland Portugal, winter is also when our model predicts the largest number of common dolphins, and this holds on the shelf, where the net fisheries operate: shelf abundance in winter was 40,575 [28,546, 56,662] against 22,826 [17,575, 29,542] in summer (Table 4). A risk assessment built on a summer density surface, the only season covered by dedicated surveys, would therefore describe the season when the fewest animals are present. A winter surface alone would not be sufficient either: per monitored day at sea, capture rates in Portuguese fisheries peak in winter for trammel nets, in summer for gillnets, in spring and summer for the beach seine and in autumn for the purse seine (DGRM, 2026b). Each fishery encounters the population at a different point of its seasonal cycle, and assessing how each fishery’s risk changes through the year requires animal density at the same seasonal resolution (Hazen et al., 2018; Maxwell et al., 2015).

Where animals concentrate is also where bycatch is recorded. Most Portuguese common dolphin strandings are recovered on the northern coast, where around three quarters of examined carcasses show evidence of bycatch (DGRM, 2026b), and the gillnet and trammel-net effort of the larger coastal vessels concentrates in the same region (Sales Henriques et al., 2024); our winter maximum lies on that northern shelf (Fig. 5A). The seasonal pattern of bycatch mortality in current ICES advice derives from strandings on the French coasts of the Bay of Biscay and western Channel (ICES, 2020, 2023b); the estimates here provide the corresponding seasonal abundance for Iberian waters, where it has been lacking. A national action plan for minimising bycatch of marine birds, mammals and reptiles is expected to enter into force in Portugal in 2026, with spatial and temporal measures for the net fisheries of this same northern coast (DGRM, 2026a). The seasonal surfaces estimated here provide the density layers against which such measures can be placed and evaluated and, overlaid with fishing effort, a basis for the plan’s risk-based monitoring.

### 4.2 Seasonal distribution, habitat and group size

The clearest ecological signal in our results is the winter maximum, which holds on the shelf and so does not rest on the extrapolated offshore surface; our mechanistic reading is confined to the shelf. Upwelling-favourable winds off western Iberia occur mainly from April to October, so upwelling-driven productivity is unlikely to explain a winter maximum: winter is dominated by downwelling, with short upwelling episodes year-round, while over the northern shelf coastal production stays relatively high through winter, sustained by river run-off and the associated low-salinity plume, a recurrent retention area for the early life stages of small pelagic fish (Relvas et al., 2007; Zwolinski et al., 2010). This is also where the winter hotspot lies (Fig. 5A). The most direct candidate is therefore the winter prey field: sardine, historically and still the principal prey of common dolphins off Portugal (Marçalo et al., 2018; Silva, 1999), spawns mainly from autumn to spring, with a broad winter peak (Coombs et al., 2006; Stratoudakis et al., 2007), almost exclusively over the shelf (Bernal et al., 2007). Consistent with this, sardine occurrence in common dolphin stomachs in adjacent Galician waters falls to its seasonal minimum in summer (Santos et al., 2013). A complementary candidate for the winter increase is redistribution within the Northeast Atlantic population, assessed and managed as a single, highly mobile unit (Murphy et al., 2021). Winter aerial surveys in the Bay of Biscay estimated lower small-delphinid point estimates than in summer (Laran et al., 2017), the opposite phase to our result, with the animals that remain concentrated over the shelf (Lambert et al., 2022; Laran et al., 2017). The pattern is consistent with a southward shift in winter, but the Biscay abundance estimates pool common dolphins with striped dolphins and no movement data demonstrates such a shift. Whether the regional gains and losses balance could be tested with a single model over the whole range built on absolute abundance (see §4.4).

Density increased in shallower, cooler and more productive water and over steeper seabed, and habitat models elsewhere in the eastern North Atlantic likewise identify cool, productive coastal waters as common dolphin habitat (Correia et al., 2019). We read these as associations with proxies for prey, not as direct drivers (Becker et al., 2014; Redfern et al., 2006). In fact, the latent field carried more structure than any single covariate (Table 3), so much of the pattern reflects processes the covariate set cannot express, plausibly the prey field, but equally anything unmodelled.

Group size varied in space, not across seasons: its seasonal means stayed between 9.4 and 10.3 animals, while its spatial variation was roughly 2.5-fold, at a finer scale than group density (posterior mean ranges 37 against 69 km, Table 3). Coupling the size field to the intensity field, and adding environmental covariates to the size model, were both tested and not supported. We report this decoupling as a finding, although the test operates at climatological resolution and cannot rule out fission–fusion processes at the scale of prey patches. The practical consequence does not depend on the mechanism though: where group size varies in space, a group-density surface can misplace animal concentrations, and the animal-density surface is the appropriate basis for management (Houldcroft et al., 2025).

### 4.3 Assumptions and limitations

As with any model, ours rests on assumptions, and statistical data integration is no universal cure for the shortcomings of the data behind it (Simmonds et al., 2020). The assumptions and limitations below are those we consider most consequential.

First, we assume certain detection on the trackline. None of the programmes carries an availability or perception correction (Laake & Borchers, 2004): no programme collected the data such a correction requires, and the few circle-back records in the Portuguese SCANS-IV component, excluded here, would not have supported estimating trackline detection on their own. Because animals are missed on the line, and predictions are made at the aerial offset, our absolute densities are minimum estimates in every season.

Second, the absolute scale depends on the choice of reference programme. We consider SCANS-IV the defensible choice, as the only randomised survey designed for abundance estimation, and we read the strongly negative cargo-vessel offset as reduced effective detectability from fast, high-sided vessels, plausibly reflecting transit speed, availability and responsive movement, rather than lower density along shipping routes.

Third, the four programmes contribute very unequally, and their effort was not placed at random. SPEA supplies 62% of the effort while SCANS-IV, the only randomised programme whose offset we use for prediction, supplies under 5% of the effort in one season. Numerical dominance of this kind is what motivates the weighting schemes proposed for integrated models, in which a large body of opportunistic records is downweighted so that it does not overwhelm a small structured dataset (Fletcher et al., 2019). We did not apply this: all four programmes are line-transect surveys with measured effort and their own detection functions, the point-process likelihood already conditions on the realised effort geometry, and any choice of weights would be subjective, as the data cannot identify them. The imbalance matters instead for prediction, and unevenly across seasons. Winter effort came almost entirely from SPEA and ATLANTIDA, both concentrated on the shelf and slope, and spring adds only a small amount of offshore CETUS effort along fixed routes. Offshore density in winter and spring is therefore a model-based extrapolation, carried by the seasonal covariate climatologies and the between-season correlation of the latent field. Restricted to the shelf, however, abundance keeps its winter maximum (Table 4), so the seasonal contrast is not an artefact of offshore extrapolation.

Fourth, group size does not enter the detection function, although large groups are more detectable than single animals. Conditioning detection on group size, as Houldcroft et al. (2025) do for a single survey, means the point process lives on an augmented space of location, distance and group size, and the likelihood integral has to be evaluated over all three. With group sizes up to 200 animals, four programmes and four seasons, the integration scheme over that augmented space becomes prohibitively large, and it was not feasible here. The observed groups are therefore a size-biased sample, and treating them as representative in the mark model likely overstates mean group size and hence animal density, partly offsetting the downward bias from assuming certain trackline detection.

Finally, pooling 2004–2026 into a seasonal climatology smooths over a dynamic ecosystem and over documented interannual variation in dolphin density in these waters (Martins et al., 2026): the surfaces are effort-weighted averages, not a description of any single year, and their credible intervals exclude interannual variance. The same smoothing applies to the covariates, which enter as seasonal climatologies (Mannocci et al., 2017); the chlorophyll-*a* effect in particular describes association with persistently productive waters rather than a response to blooms. Therefore, we present the surfaces as climatological baselines rather than estimates for any particular year.

### 4.4 What data integration contributed, and outlook

The clearest contribution of integration is spatiotemporal coverage. Dedicated surveys are expensive and logistically demanding, which is why line-transect data for cetaceans are sparse. Opportunistic programmes that observe from platforms already at sea, or that run year-round for other purposes, accumulate effort at a fraction of the cost. Integration puts them together, and the effort collected outside any dedicated design fills the seasons, years and areas a decadal summer survey cannot reach. The one-stage formulation propagates detection, field, group-size and process uncertainty into abundance within a single likelihood, avoiding the separate variance propagation that two-stage density surface methods require. Another advantage of our model lies in the point-process formulation: sightings cluster beyond what the covariates explain, and the latent field of the LGCP models this spatial autocorrelation directly; ignoring it would overstate the precision of the estimates (Illian et al., 2008). In addition, any data stream whose observation process can be written as a thinning of the shared intensity can enter through its own observation model, for example presence-only sightings (Renner et al., 2015) and passive acoustic detections (Schliep et al., 2024).

Several extensions follow naturally. The most important is incorporating trackline-detection data, such as circle-back records or double-observer effort, into the joint likelihood. This will allow to relax the certain-detection assumption and move the absolute scale from a minimum towards an unbiased estimate. The building blocks exist: mark–recapture distance sampling supplies the estimators (Laake & Borchers, 2004), Bayesian formulations of distance sampling are established (Oedekoven et al., 2014), including mark–recapture distance sampling (Sigourney et al., 2020), but not yet, to our knowledge, within the point-process formulation. Further, the temporal structure can be refined: year-resolved or finer-than-seasonal fields would recover the variation the climatology smooths over and separate trend from average pattern, though estimating a spatial field at a finer temporal resolution than per season carries a high computational cost. Finally, the model domain could be extended. Seasonal line-transect data exist for the Bay of Biscay, the English Channel and further north, and nothing prevents a model built on all available data across European Atlantic waters, which could test whether the winter increase off Portugal corresponds to a decrease elsewhere in the range.

Seasonal closures, bycatch risk assessment and status reporting all require knowing how many animals are where, in each season. For common dolphins off mainland Portugal this information did not exist: dedicated surveys provided summer snapshots, and opportunistic programmes seasonal coverage without a design-backed absolute scale. The OSPAR Quality Status Report 2023 links its low data availability for cetaceans to gaps in coverage offshore and in winter, and calls for better use and coordination of surveys beyond the dedicated programmes, alongside more frequent dedicated surveys (Geelhoed et al., 2022). Integrated models are no substitute for dedicated surveys: the absolute scale of this analysis rests on SCANS-IV, and each new dedicated survey improves it. The summer surface was checked against design-based estimates (Appendix S2), but no design-based counterpart exists for the other seasons, and the winter and spring surfaces rest partly on model-based extrapolation offshore (§4.3). Dedicated effort outside summer, repeated across seasons and reaching offshore in winter, would provide the first independent test of the seasonal pattern estimated here. Integration, meanwhile, increases what is learned from all other effort, and since every programme is described by its own observation model, the same construction can be applied to any wildlife population monitored through several partial programmes (Isaac et al., 2020).

## Acknowledgements

The project that gave rise to these results received the support of a fellowship from the ‘la Caixa’ Foundation (ID 100010434) to Moritz Klaassen. The fellowship code is LCF/BQ/DI23/11990054. FCT – Fundação para a Ciência e a Tecnologia, I.P. supported Filipe Alves and Marc Fernandez through the strategic projects UID/04292/2025 (https://doi.org/10.54499/UID/04292/2025), granted to MARE, and LA/P/0069/2020 (https://doi.org/10.54499/LA/P/0069/2020), granted to ARNET; Filipe Alves through a research contract (https://doi.org/10.54499/2023.08949. CEECIND/CP2851/CT0001); Tiago A. Marques through CEAUL, UID/00006/2025 (https://doi.org/10.54499/UID/00006/2025); Miguel P. Martins through PhD grant 2025.04354.BD; and Ana M. Correia through contract 2024.08759.CEECIND. Tiago A. Marques was also funded by the European Union – NextGenerationEU through UID/PRR/00006/2025 (https://doi.org/10.54499/UID/PRR/00006/2025) and by the AMERICANO project, and Marc Fernandez by the Atlantic Whale Deal project (EAPA_0004/2022), co-financed by Interreg Atlantic Area 2021–2027. Andrew Houldcroft was funded by a doctoral training grant awarded through the UKRI AI Centre for Doctoral Training in Environmental Intelligence (EP/S022074/1). The contributions of Ana M. Correia and Iolanda M. Silva were funded by the CIIMAR monitoring programme, the CETUS Project and ATLANTIDA II (NORTE2030-FEDER-01799200; NORTE 2030, ERDF and FCT), and by the European Commission’s Recovery and Resilience Facility through UID/04423/2025 (https://doi.org/10.54499/UID/04423/2025), UID/PRR/04423/2025 (https://doi.org/10.54499/UID/PRR/04423/2025) and LA/P/0101/2020 (https://doi.org/10.54499/LA/P/0101/2020). CCMAR received FCT funding through UID/04326/2025 (https://doi.org/10.54499/UID/04326/2025), UID/PRR/04326/2025 (https://doi.org/10.54499/U ID/PRR/04326/2025) and LA/P/0101/2020 (https://doi.org/10.54499/LA/P/0101/2020). We thank Anita Gilles for access to the SCANS-IV data. Portuguese SCANS-IV data collection was coordinated by Catarina Eira and carried out by the University of Aveiro, funded by the Fundo Ambiental of the Portuguese Ministry of the Environment and by CESAM through FCT (UID/50017/2025, https://doi.org/10.54499/UID/50017/2025; LA/P/0094/2020, https://doi.org/10.54499/LA/P/0094/2020).

## Author Contributions

Moritz Klaassen, Tiago A. Marques, Marc Fernandez and Filipe Alves conceived the study. Moritz Klaassen designed and implemented the model, performed the analyses, prepared the figures and led the writing. Finn Lindgren and Sara Martino advised on the statistical model, Andrew Houldcroft on the group-size methodology, Miguel P. Martins on the analysis and interpretation, and Ana Marçalo on the bycatch management context. Ana M. Correia and Iolanda M. Silva curated and provided the CETUS and ATLANTIDA data, Nuno Oliveira curated and provided the SPEA data, and Andreia Torres-Pereira provided access to the Portuguese SCANS-IV data. Tiago A. Marques, Marc Fernandez and Filipe Alves supervised the work. All authors contributed critically to the drafts and gave final approval for publication.

## Supporting Information

### This file contains

- Appendix S1. Supplementary methods and results
- Appendix S2. Comparison with previous estimates
- Appendix S3. Mean and variance of the number of animals

## Appendix S1. Supplementary methods and results

### S1.1 Environmental covariates

Table S1 lists the eleven candidate covariates and their sources, and Figure S1 their pairwise Pearson correlations on the prediction grid for each seasonal climatology. Collinearity was assessed on each seasonal climatology separately, on a random sample of 10,000 grid cells, using pairwise correlations and variance inflation factors (VIF). Among the eleven candidates, VIFs exceeded 10 in at least one season for sea surface temperature (10.7–29.6 across seasons), zooplankton (15.9–32.6), mixed-layer depth (6.7–18.5), salinity (8.3–22.3) and chlorophyll-*a* (2.9–11.7). Salinity was strongly correlated with sea surface temperature in every season (*r* = 0.75–0.95); mixed-layer depth with bathymetry (*r* = 0.72–0.89) and, in spring and autumn, with distance to coast (*r* = 0.76 and 0.80); and zooplankton with chlorophyll-*a* (*r* = 0.73– 0.83) and with sea surface temperature (*r* = −0.75 to −0.87). Salinity, mixed-layer depth and zooplankton were therefore removed, retaining in each case the variable with the more direct ecological interpretation. Among the eight remaining covariates the strongest association was between bathymetry and distance to coast (*r* = 0.69 in all seasons); the largest VIF was 5.2 (distance to coast) and all others were at most 3.1. These eight entered the backward elimination described in Section S1.4.

**Table S1:** Candidate environmental covariates for the group density model. Seasonal covariates are climatological means for each of the four seasons over 2004–2026.

| Covariate | Unit | Source | Temporal |
| --- | --- | --- | --- |
| Sea surface temperature | °C | CMEMS IBI physics reanalysis | Seasonal |
| Sea surface salinity | PSU | CMEMS IBI physics reanalysis | Seasonal |
| Mixed-layer depth | m | CMEMS IBI physics reanalysis | Seasonal |
| Bathymetry | m | CMEMS IBI physics reanalysis | Static |
| Chlorophyll-*a* | mg m<sup>-3</sup> | CMEMS Atlantic ocean colour | Seasonal |
| Zooplankton | mmol C m<sup>-3</sup> | CMEMS IBI biogeochemistry | Seasonal |
| Sea surface temperature gradient | °C km<sup>-1</sup> | Derived (sea surface temperature) | Seasonal |
| Seabed slope | degrees | Derived (bathymetry) | Static |
| Distance to coast | km | Derived (Natural Earth land polygons) | Static |
| Distance to canyon | km | Derived (EMODnet geomorphology) | Static |
| Distance to seamount | km | Derived (EMODnet geomorphology) | Static |

**Figure S1.**
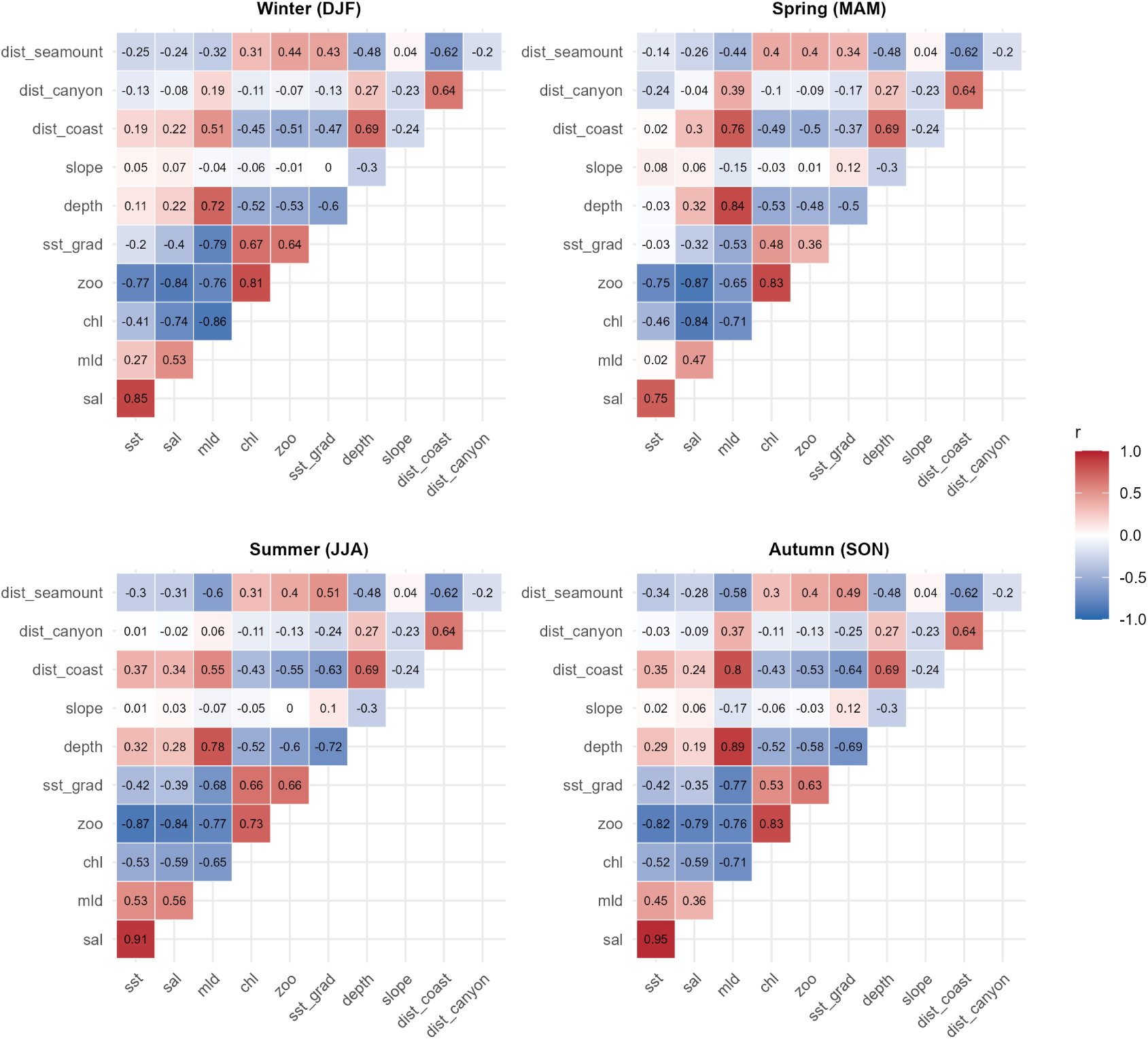
Pairwise Pearson correlations among the eleven candidate covariates on the prediction grid, by season.

### S1.2 Survey data and detection functions

Detection covariates were limited to those recorded on effort as well as at sightings, because the point-process likelihood evaluates the detection function over the whole sampler. The candidate sets were sea state for SPEA; sea state, platform height, and the most experienced (MEO) and cumulative (CUM) observer experience scores for CETUS; sea state, wind state and visibility for ATLANTIDA; and sea state, glare and cloud cover for SCANS-IV. For each programme, the half-normal and hazard-rate key functions were fitted with each candidate covariate in turn and without covariates, and compared by DIC. Continuous covariates were standardised and entered linearly on the log scale of the detection scale parameter; categorical covariates entered as factor contrasts. The retained model for each programme are given below.

#### SPEA

Two series collected under the same European Seabirds at Sea protocol were combined: the 2004–2020 campaigns analysed by Martins et al. (2026) and new campaigns from 2021–2026. Effort is concentrated on the shelf along the west and south coasts in all seasons, with offshore transects at much lower intensity (Figure S2A). Perpendicular distance was recorded in four bands (0–50, 50–100, 100–200 and 200–300 m). The outer band edge, 300 m, is the truncation distance, so no sightings were discarded (Figure S2B). Sea state entered the detection function as a factor with levels 0, 1, 2, 3 and 4 or more.

**Figure S2.**
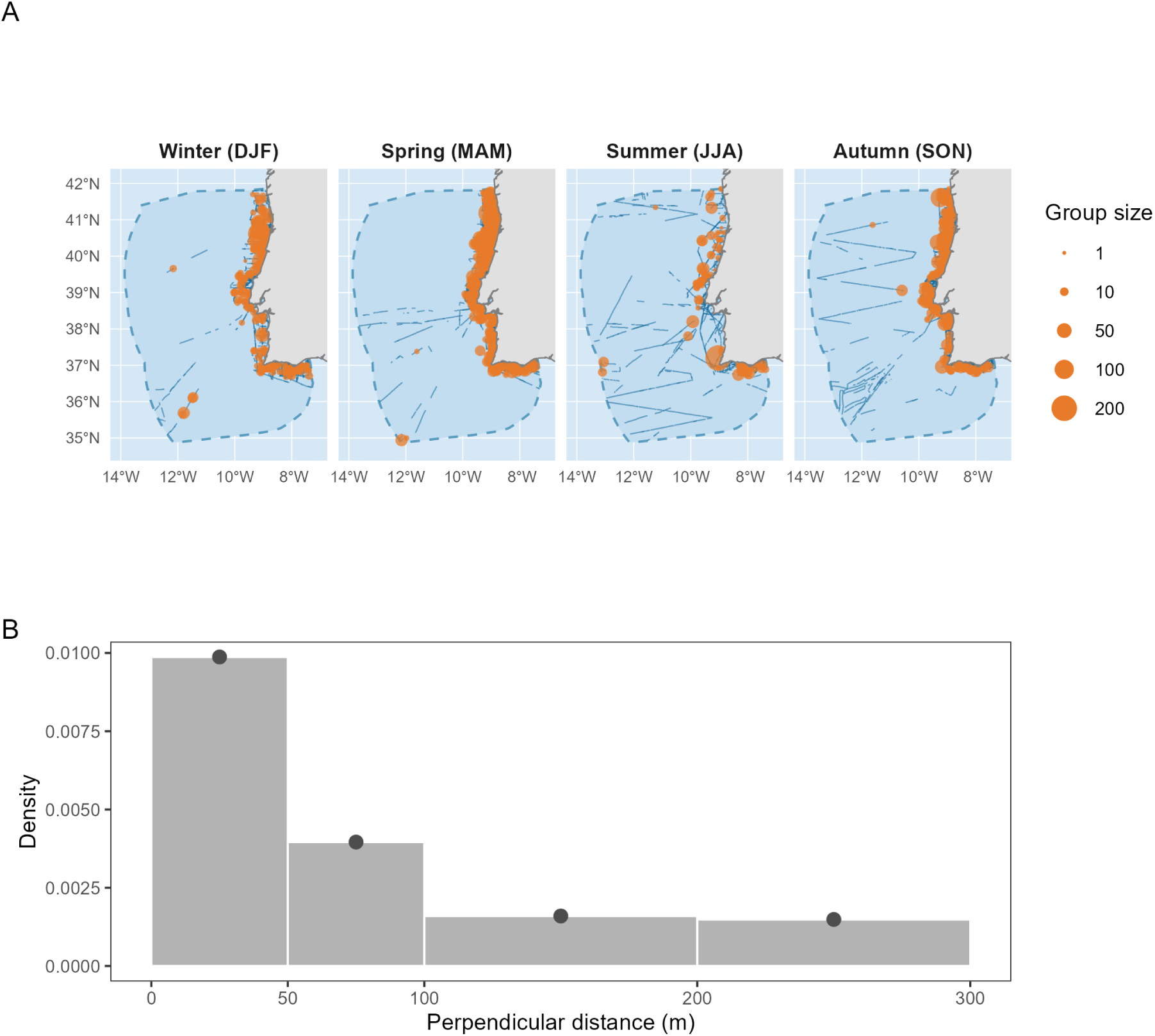
SPEA, 2004–2026. (A) On-effort transects (blue) and common dolphin sightings (orange, scaled by group size) by season within the model domain (dashed outline). (B) Perpendicular distance distribution of sightings.

**Table S2:** SPEA effort and common dolphin sightings by season, 2004–2026. ER, encounter rate.

| Season | Effort (km) | Groups | Animals | ER (groups/100 km) | ER (animals/100 km) |
| --- | --- | --- | --- | --- | --- |
| Winter (DJF) | 14,557 | 237 | 2,172 | 1.63 | 14.9 |
| Spring (MAM) | 47,920 | 482 | 3,876 | 1.01 | 8.1 |
| Summer (JJA) | 13,346 | 99 | 901 | 0.74 | 6.8 |
| Autumn (SON) | 24,785 | 359 | 3,267 | 1.45 | 13.2 |

The hazard-rate with sea state had the lowest DIC (Table S3), but the half-normal with sea state was retained. With distances in four bands the detection function is informed by four points only, and the hazard-rate shape parameter is poorly constrained. The half-normal gives a smooth decline consistent with the band counts, and the same choice was made for this protocol by Martins et al. (2026). Sea state was retained in either case.

**Table S3:**
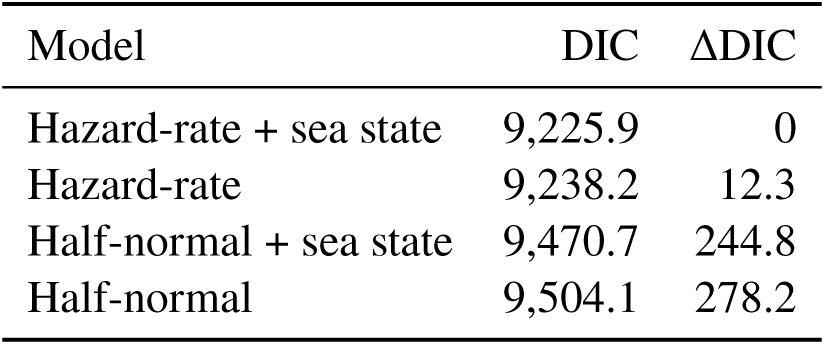
SPEA detection function comparison.

#### CETUS

Cargo and oceanographic vessel transects follow fixed commercial routes between mainland Portugal and the Atlantic islands, so effort reaches far offshore but along a few repeated lines (Figure S3A). Perpendicular distances were truncated at 1,000 m, removing 5 of 339 sightings (Figure S3C).

**Figure S3.**
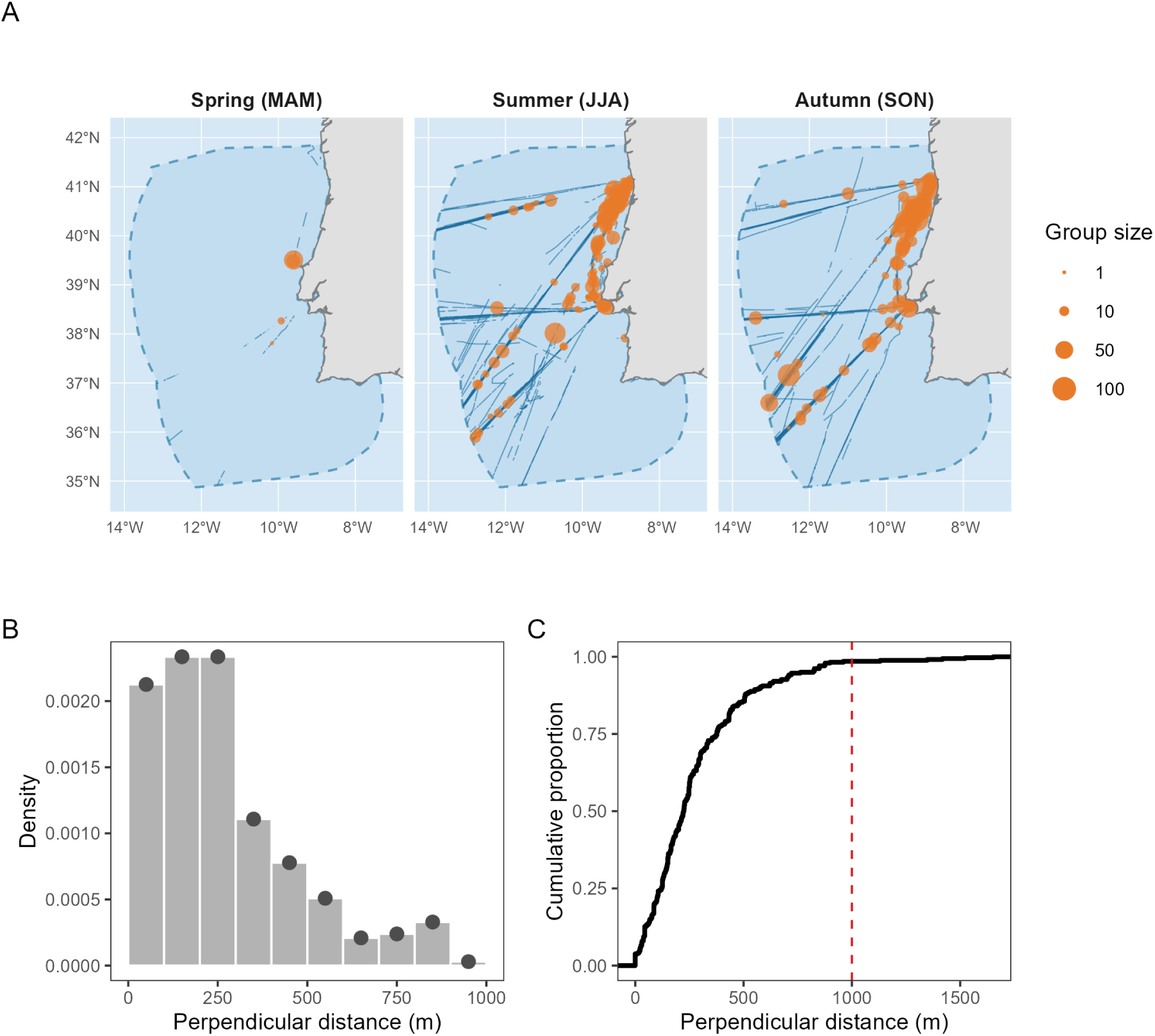
CETUS, 2012–2024. (A) On-effort transects (blue) and common dolphin sightings (orange, scaled by group size) by season within the model domain (dashed outline). (B) Perpendicular distance distribution of sightings within the truncation distance. (C) Cumulative distribution of all recorded distances; the dashed line marks the truncation distance.

**Table S4:**
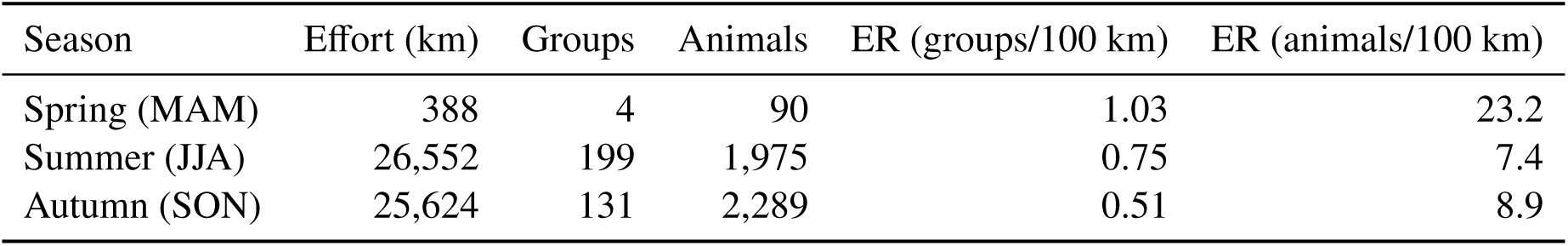
CETUS effort and common dolphin sightings by season, 2012–2024, after truncation at 1,000 m.

| Season | Effort (km) | Groups | Animals | ER (groups/100 km) | ER (animals/100 km) |
| --- | --- | --- | --- | --- | --- |
| Spring (MAM) | 388 | 4 | 90 | 1.03 | 23.2 |
| Summer (JJA) | 26,552 | 199 | 1,975 | 0.75 | 7.4 |
| Autumn (SON) | 25,624 | 131 | 2,289 | 0.51 | 8.9 |

Each CETUS observer receives an experience score from 0 to 20, based on experience at sea, identification, range of species and protocol. With two observers on effort, we used the cumulative score of both (0–40), standardised before fitting, as a continuous covariate on the detection scale, and tested the score of the most experienced observer (MEO) as an alternative (Table S5). The hazard-rate was clearly preferred over the half-normal, but the five hazard-rate models were indistinguishable by DIC, all within 1.7 units. We therefore chose on external grounds and retained the cumulative observer-experience score, the observer-level bias documented for this programme by Oliveira-Rodrigues et al. (2022) on the full CETUS dataset, of which the sightings used here are a subset.

**Table S5:** CETUS detection function comparison. MEO, score of the most experienced observer; CUM, cumulative score of both observers.

| Model | DIC | ΔDIC |
| --- | --- | --- |
| Hazard-rate + sea state | 3,787.6 | 0 |
| Hazard-rate | 3,788.1 | 0.5 |
| Hazard-rate + MEO | 3,788.2 | 0.6 |
| Hazard-rate + CUM | 3,789.2 | 1.6 |
| Hazard-rate + platform height | 3,789.3 | 1.7 |
| Half-normal + MEO | 3,795.9 | 8.3 |
| Half-normal + sea state | 3,796.6 | 9.0 |
| Half-normal + CUM | 3,796.8 | 9.2 |
| Half-normal | 3,797.5 | 9.9 |
| Half-normal + platform height | 3,799.6 | 12.0 |

#### ATLANTIDA

Only the campaigns following standardised parallel transects off the northern coast (Figure S4A) were used; the campaigns near Porto are non-systematic searches without distance measurement and were excluded. Perpendicular distances were truncated at 500 m, removing 2 of 69 sightings (Figure S4C).

**Figure S4.**
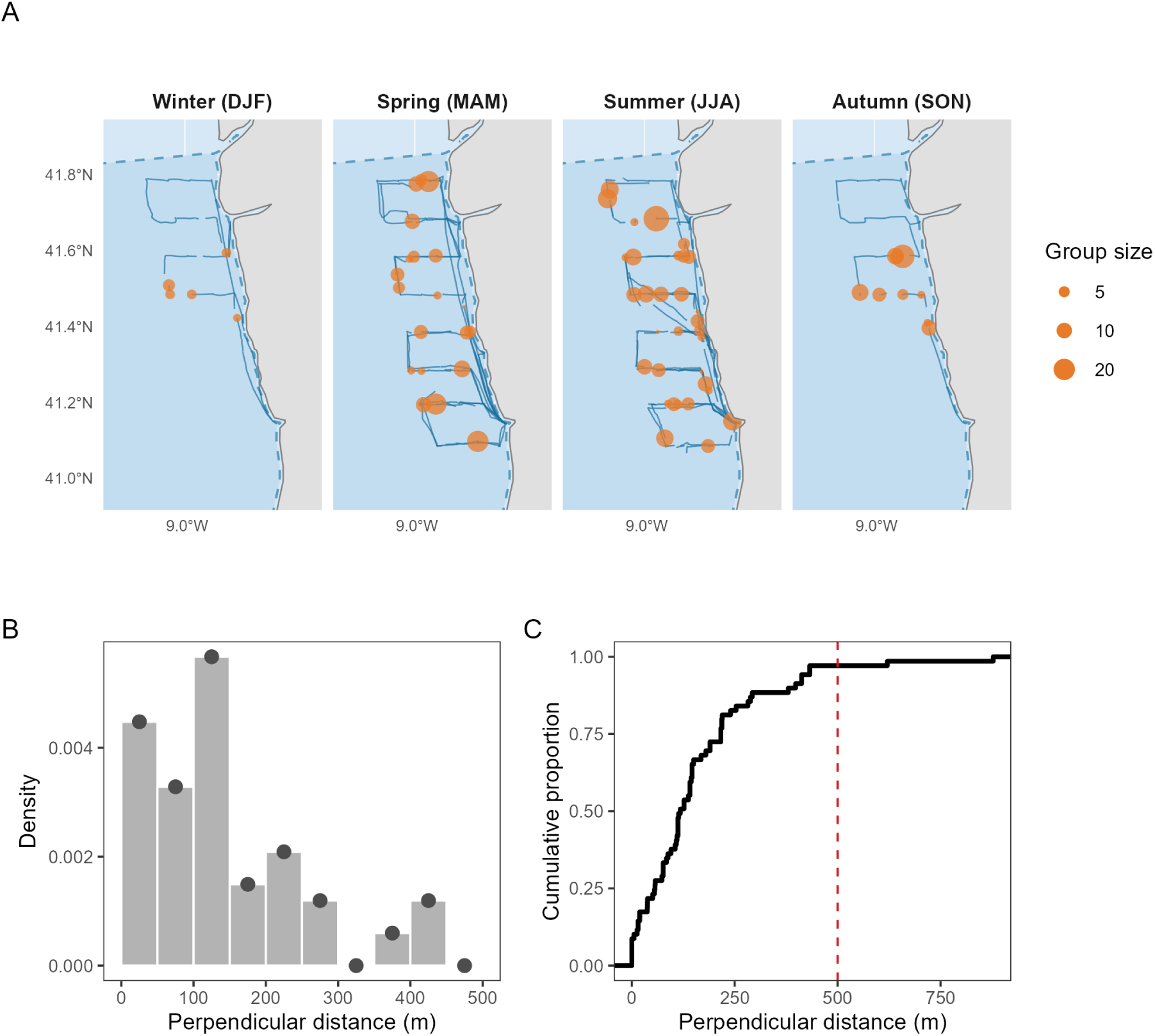
ATLANTIDA, 2021–2024. (A) On-effort transects (blue) and common dolphin sightings (orange, scaled by group size) by season off the northern Portuguese coast; the dashed line is the model domain boundary. (B) Perpendicular distance distribution of sightings within the truncation distance. (C) Cumulative distribution of all recorded distances; the dashed line marks the truncation distance.

With 67 sightings spread over several levels of wind state, sea state and visibility, individual factor contrasts rested on fewer than ten sightings and could not be estimated with any precision, so no detection covariate was used. The hazard-rate was retained.

**Table S6:**
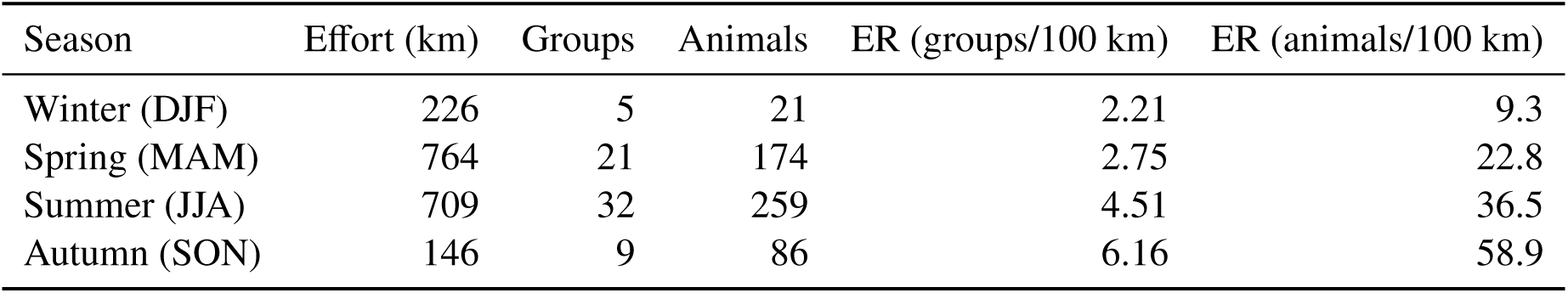
ATLANTIDA effort and common dolphin sightings by season, 2021–2024, after truncation at 500 m.

#### SCANS-IV

The Portuguese blocks of the SCANS-IV aerial survey were flown in July and August 2022 on equal-spaced zigzag and parallel designs, giving the only spatially balanced coverage of the offshore domain (Figure S5A). Records from circle-back procedures were excluded. Perpendicular distances were truncated at 400 m, removing 5 of 183 sightings (Figure S5C).

**Figure S5.**
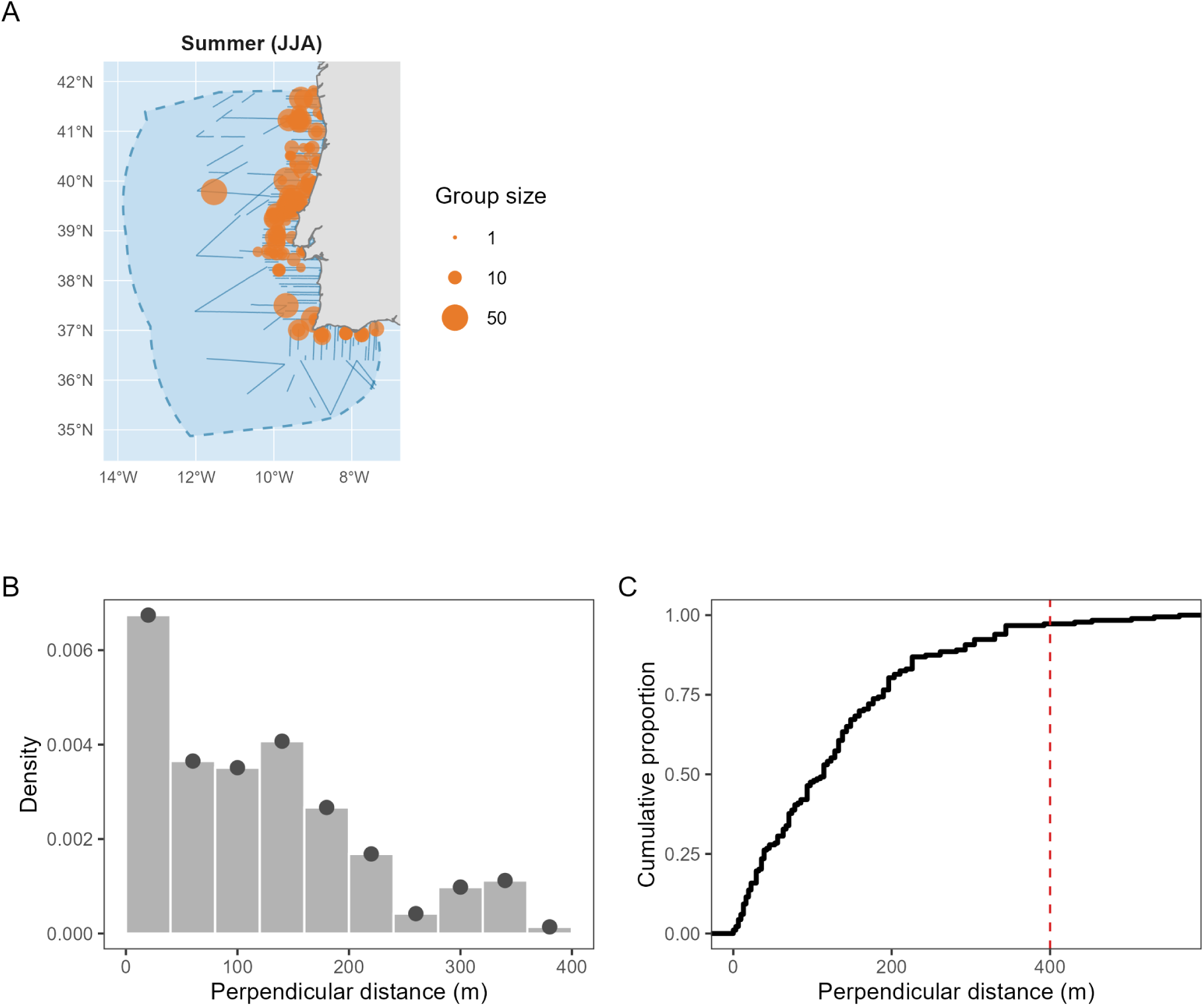
SCANS-IV, summer 2022. (A) On-effort transects (blue) and common dolphin sightings (orange, scaled by group size) within the model domain (dashed outline). (B) Perpendicular distance distribution of sightings within the truncation distance. (C) Cumulative distribution of all recorded distances; the dashed line marks the truncation distance.

**Table S7:** SCANS-IV effort and common dolphin sightings, summer 2022, after truncation at 400 m.

| Season | Effort (km) | Groups | Animals | ER (groups/100 km) | ER (animals/100 km) |
| --- | --- | --- | --- | --- | --- |
| Summer (JJA) | 7,498 | 178 | 1,890 | 2.37 | 25.2 |

Glare and sea state gave the lowest DICs, within 0.6 units of each other, and their half-normal versions within 1.2 (Table S8). Sea state was retained: its effect was monotonic and interpretable, whereas the glare contrasts showed no consistent ordering across levels. Cloud cover was not supported.

**Table S8:** SCANS-IV detection function comparison.

| Model | DIC | ΔDIC |
| --- | --- | --- |
| Hazard-rate + glare | 1,267.5 | 0 |
| Hazard-rate + sea state | 1,268.1 | 0.6 |
| Half-normal + sea state | 1,268.5 | 1.0 |
| Half-normal + glare | 1,268.7 | 1.2 |
| Half-normal | 1,274.7 | 7.2 |
| Half-normal + cloud cover | 1,277.3 | 9.8 |
| Hazard-rate | 1,277.3 | 9.8 |
| Hazard-rate + cloud cover | 1,280.4 | 12.9 |

### S1.3 Mesh construction and integration

The latent fields *ω* and *φ* are Gaussian random fields with Matérn covariance. For two points separated by Euclidean distance *r* = || **s** − **s**^′^||,

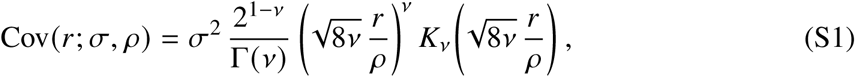

with *K_ν_* the modified Bessel function of the second kind, *σ* the marginal standard deviation and *ρ* the practical range. The smoothness *ν* is fixed at 1, as is standard for two-dimensional applications. Following Lindgren et al. (2011), each field is represented as the solution of a stochastic partial differential equation, discretised on the finite-element mesh described below, which yields a sparse Gaussian Markov random field. All spatial computation was carried out in a Lambert azimuthal equal-area projection centred at 38.5^◦^N, 10.5^◦^W, in kilometres. The model domain is the mainland Portuguese EEZ with land removed, and it is the domain over which density was predicted and abundance integrated. The finite-element mesh for the SPDE representation of the two Matérn fields (Figure S6) was built with the fmesher package. Node locations were set explicitly rather than by triangle-size constraints: a hexagonal lattice of 15 km spacing covered the domain buffered by 30 km, and a second lattice of 7.5 km spacing was superimposed within 60 km of the coast, where survey effort and sightings are concentrated. The mesh extends beyond the domain to a non-convex hull 150 km wide, triangulated coarsely, so that the boundary effects inherent to the SPDE approximation fall outside the region of interest.

The integral in the point-process likelihood (Eq. 7 of the main text) was evaluated numerically over the product of space, perpendicular distance and season.

**Figure S6.**
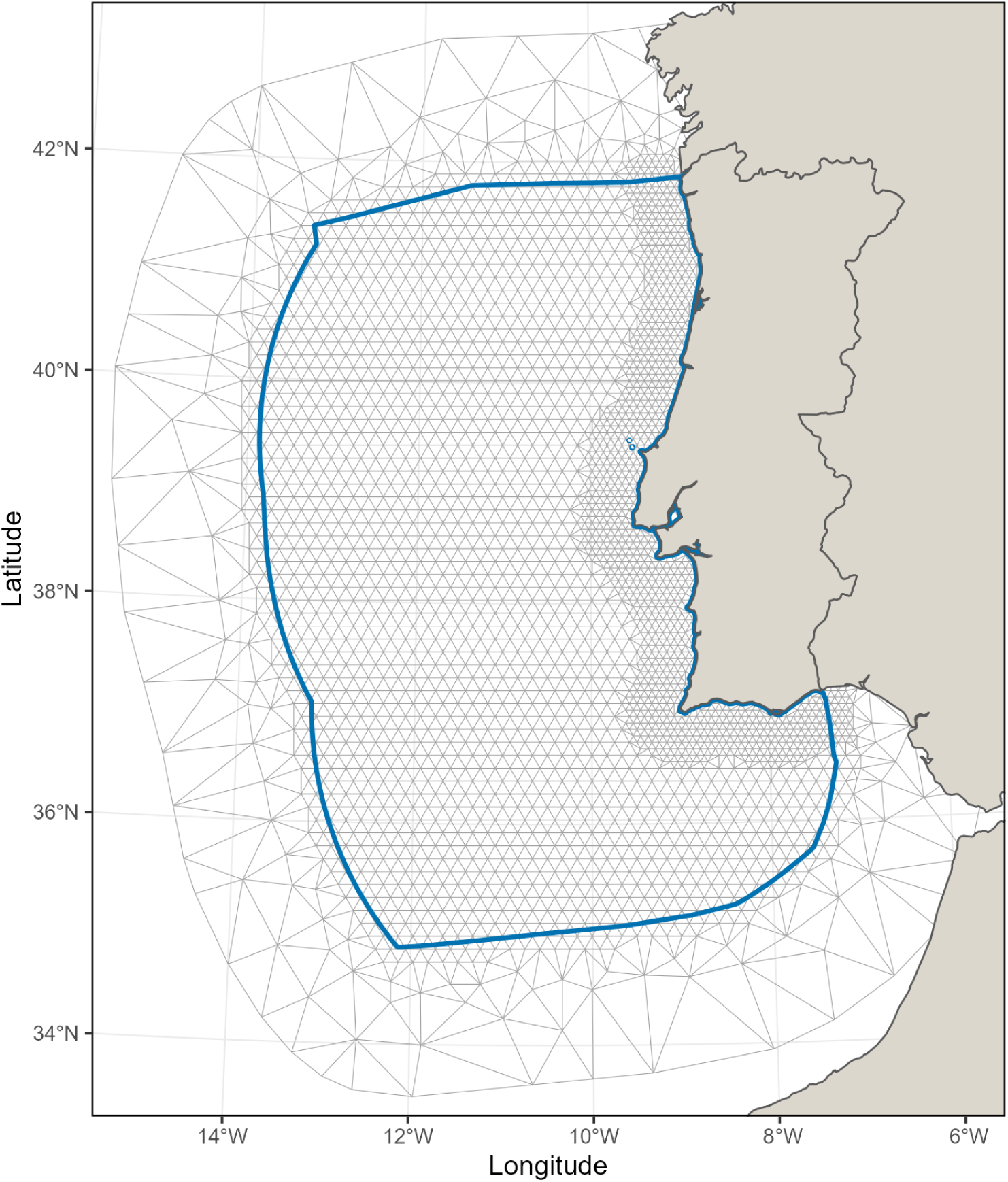
Finite-element mesh used for the SPDE approximation to the Matérn field. Grey lines show the mesh triangulation; the blue outline delimits the Portuguese mainland Exclusive Economic Zone (EEZ) excluding land, which is the domain over which density was predicted and abundance integrated.

### S1.4 Model selection

The covariates of the group density model were selected by backward elimination on DIC, starting from the model with all eight candidates retained after collinearity screening (Section S1.1). Every candidate model kept the full structure of the final model; only the linear covariate terms in the intensity changed. At each step every covariate in the current set was removed in turn, the removal giving the lowest DIC was accepted if it improved on the current model, and elimination stopped otherwise.

The sea-surface-temperature gradient was removed first, then distance to seamount, then distance to coast; at the fourth step no removal improved DIC and elimination stopped (Table S9). The retained set is sea surface temperature, chlorophyll-*a*, bathymetry, seabed slope and distance to canyon.

**Table S9:** Backward elimination of intensity covariates by DIC. Each cell is the DIC of the model obtained by removing the row covariate from the set current at that step; the current model’s DIC is in the header row. The lowest DIC at each step is in bold and that covariate was removed if it improved on the current model. Dashes mark covariates already removed.

|  | Step 1 | Step 2 | Step 3 | Step 4 |
| --- | --- | --- | --- | --- |
| Current model DIC | 24,027.0 | 24,025.2 | 24,024.1 | 24,023.2 |
| Sea surface temperature | 24,037.7 | 24,036.5 | 24,036.0 | 24,034.8 |
| Chlorophyll-*a* | 24,040.6 | 24,039.4 | 24,038.6 | 24,037.7 |
| Bathymetry | 24,032.1 | 24,029.9 | 24,029.9 | 24,029.7 |
| Seabed slope | 24,040.6 | 24,038.9 | 24,038.3 | 24,038.3 |
| Distance to canyon | 24,027.7 | 24,025.6 | 24,025.2 | **24,023.6** |
| Distance to coast | 24,026.0 | 24,024.3 | **24,023.2** | — |
| Distance to seamount | 24,026.4 | **24,024.1** | — | — |
| Sea surface temperature gradient | **24,025.2** | — | — | — |

## Appendix S2. Comparison with previous estimates

This appendix compares predictions from the fitted model with previously published estimates for the same waters. The four SCANS-IV blocks with common dolphin sightings lie entirely within our study area, so abundance can be compared directly; the SCANS-III blocks AA and AB extend beyond it, so mean densities are compared over the overlap between each block and the study area only. SCANS estimates are design-based and corrected for detection on the trackline, whereas ours assume certain detection and are minimum estimates. Estimates of Martins et al. (2026) are model-based, derived from the 2004–2020 SPEA data, a subset of the data integrated here; those rows are therefore a consistency check between models rather than a validation against independent data.

The only design-based estimates of common dolphin density for these waters come from the SCANS surveys, and our summer surface can be checked against them at the scale of their survey blocks. Predicted summer abundance fell within the published 95% confidence intervals in all four SCANS-IV blocks with common dolphin sightings, with posterior means below the design-based estimates in each, as the certain-detection assumption implies, and closest in the block that held most of the sightings (IC-G: 39,988 (31,308–50,235) against 43,843 (26,812–75,330); Table S10). Because the SCANS-IV sightings enter our model, this is a comparison of estimators on shared data rather than an independent validation; the agreement shows that integrating three further programmes did not pull the summer surface away from the design-based estimates. The independent check is SCANS-III, whose data do not enter the model: predicted densities in the two blocks overlapping our study area were roughly half the corrected 2016 values. This shortfall is consistent in direction and size with uncorrected trackline detection, but interannual variability contributes too: the comparison sets a multi-year climatology against a single summer, in waters where dolphin density is documented to vary between years (Martins et al., 2026), and the two contributions cannot be separated.

**Table S10:** Predicted abundance and density (posterior mean, 95% credible interval) against previously published estimates. SCANS values are design-based and corrected for trackline detection, with 95% confidence intervals; the SCANS-III intervals were reconstructed from the reported coefficients of variation assuming a lognormal distribution. SCANS-III rows compare mean densities (individuals per km^2^) over the study-area overlap only. Martins et al. (2026) values are the median and range of their annual estimates (2004–2020), not confidence intervals. For the EEZ domain, which is the full study area, our values are the abundance estimates reported in the main text. Areas are those of the compared domains.

|  | Area (km<sup>2</sup>) | This study | Previous estimate |
| --- | --- | --- | --- |
| *SCANS-IV blocks, summer 2022 (abundance)* |  |  |  |
| IC-C | 15,708 | 6,621 ( 4,305–10,112) | 14,367 ( 4,556–29,900) |
| IC-E | 44,901 | 5,169 ( 2,966– 8,561) | 8,548 ( 50–39,839) |
| IC-F | 51,017 | 7,561 ( 4,323–12,635) | 9,857 ( 98–28,905) |
| IC-G | 42,131 | 39,988 (31,308–50,235) | 43,843 (26,812–75,330) |
| *SCANS-III blocks, summer 2016 (density, overlap only)* |  |  |  |
| AA | 3,794 | 0.773 (0.496–1.183) | 1.536 (0.484–4.873) |
| AB | 19,443 | 1.116 (0.849–1.439) | 2.372 (1.402–4.012) |
| *Martins et al. (2026) coastal band, DTC < 30 km (abundance)* |  |  |  |
| Winter | 24,315 | 38,437 (27,496–53,599) | 21,492 (18,713–29,664) |
| Spring | 24,315 | 19,559 (14,490–26,336) | 17,249 (13,737–25,887) |
| Summer | 24,315 | 24,572 (18,691–31,876) | 29,318 (23,843–34,340) |
| Autumn | 24,315 | 31,324 (23,352–41,351) | 22,468 (19,052–34,179) |
| *Martins et al. (2026) EEZ domain (abundance)* |  |  |  |
| Summer | 314,130 | 76,908 (59,505–97,901) | 95,026 (62,237–108,640) |
| Autumn | 314,130 | 86,435 (64,301–114,451) | 169,421 (97,635–194,762) |

**Figure S7.**
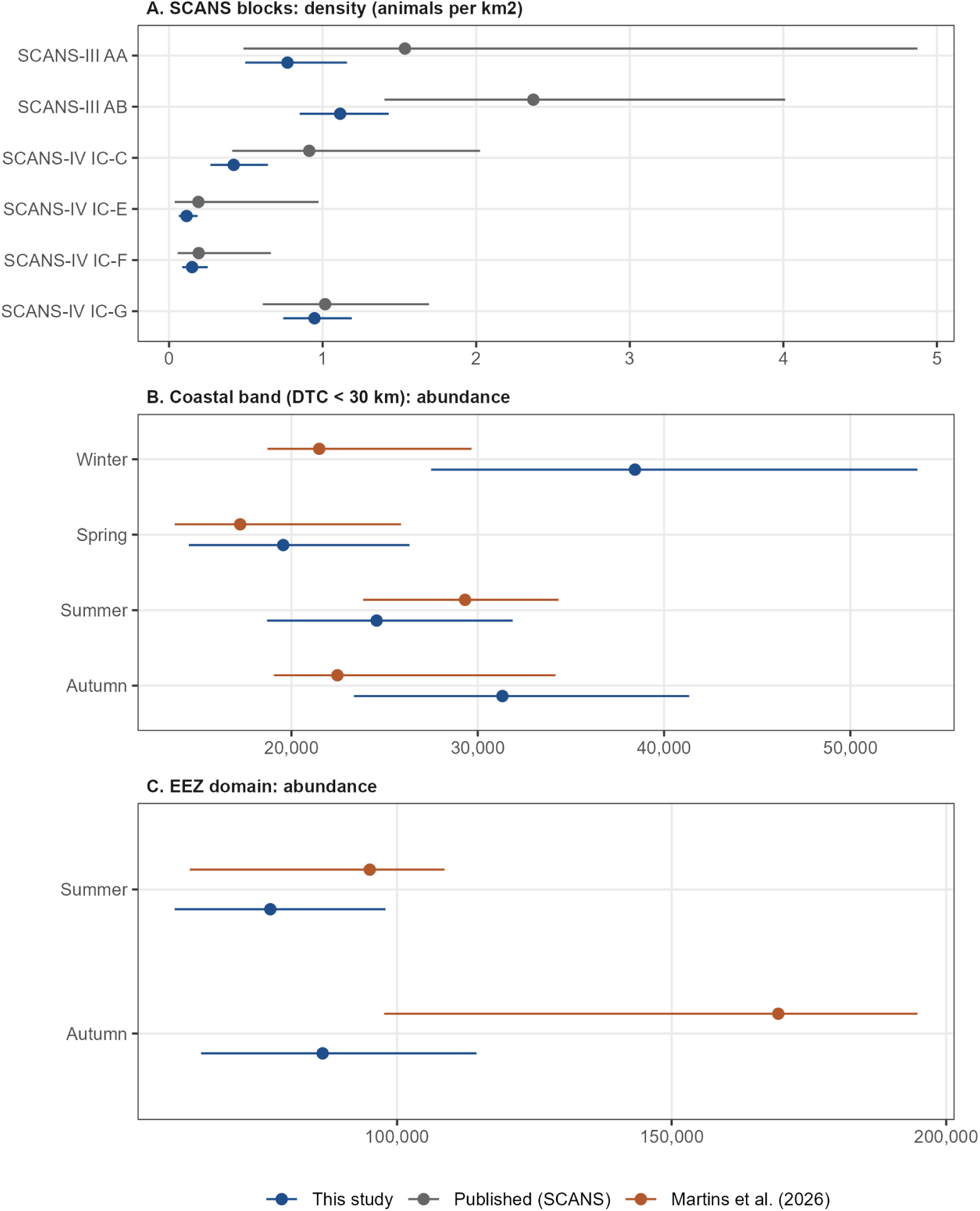
Predictions of this study (posterior mean and 95% credible interval) against previous estimates, for each comparison domain in Table S10. The intervals differ in meaning by source: credible intervals for this study; 95% confidence intervals for the SCANS estimates, reconstructed from the reported coefficients of variation for SCANS-III; and the range of annual estimates (2004–2020) for Martins et al. (2026).

The only other seasonal estimates for these waters are the model-based ones of Martins et al. (2026). Predicted onto their reporting domains (Figure S10), the two models agree well where the shared data are dense: in the coastal band, our spring and summer estimates fall within the range of their annual estimates, and both models place the coastal minimum in spring. The two divergences have different sources. In winter in the coastal band, our 38,437 (27,496–53,599) is nearly twice their median of 21,492 (annual range 18,713–29,664): our estimate draws on six further years of winter effort, the catamaran programme, and the between-season correlation of the latent field, which allows the better-surveyed seasons to inform winter and overall density to differ between seasons in ways their environmental covariates alone cannot capture. In autumn over the EEZ, their 169,421 is roughly twice our 86,435 (64,301–114,451): their estimate rests on a seasonally varying distance-to-coast effect carried into offshore waters with little SPEA effort, with an uncertainty they themselves note limits interpretation, while roughly 20,000 km of autumn cargo-vessel effort in oceanic waters holds our offshore surface low.

**Figure S8.**
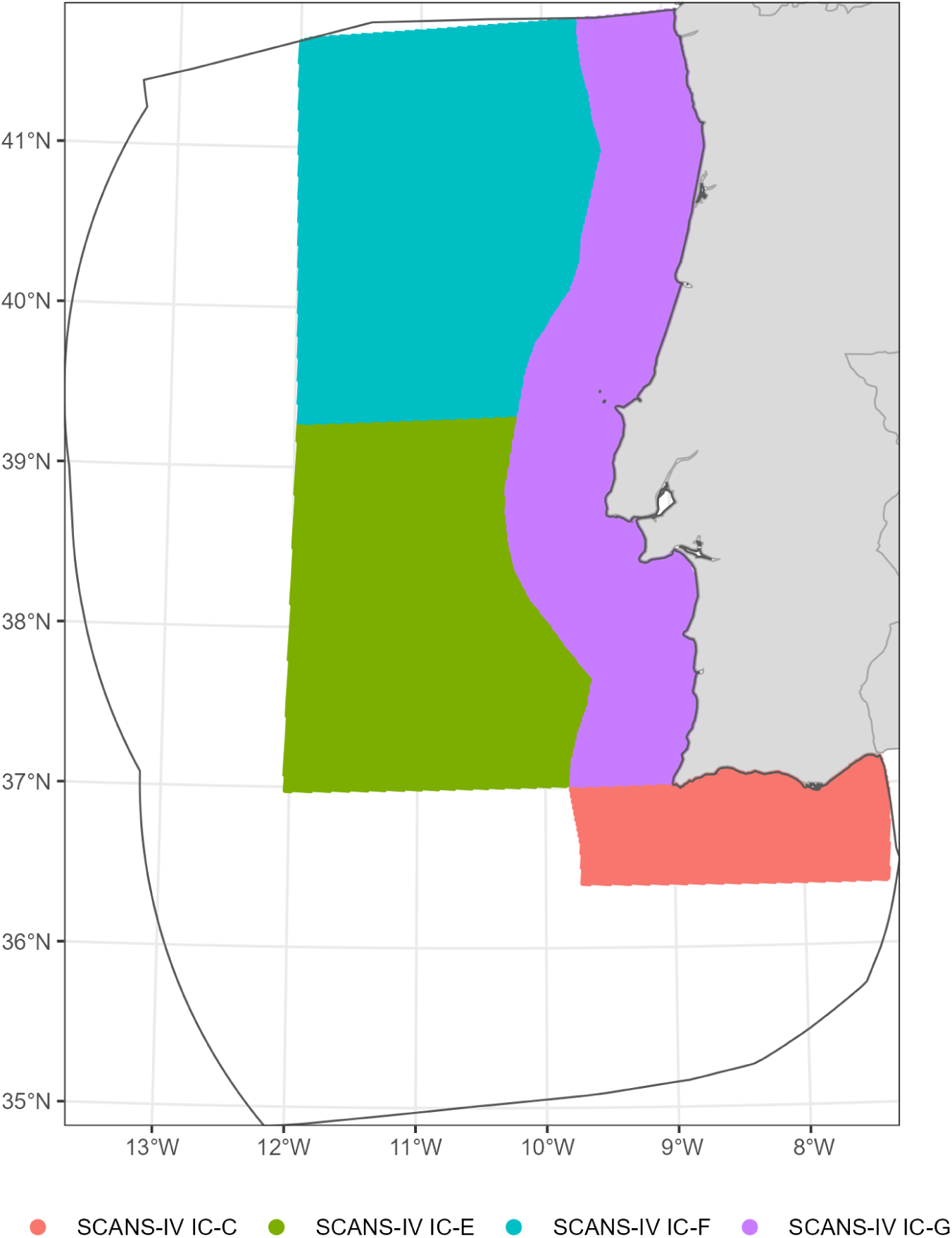
The four SCANS-IV survey blocks with common dolphin sightings (IC-C, IC-E, IC-F, IC-G) and the prediction cells entering each block estimate in Table S10. All four blocks lie within the study area.

**Figure S9.**
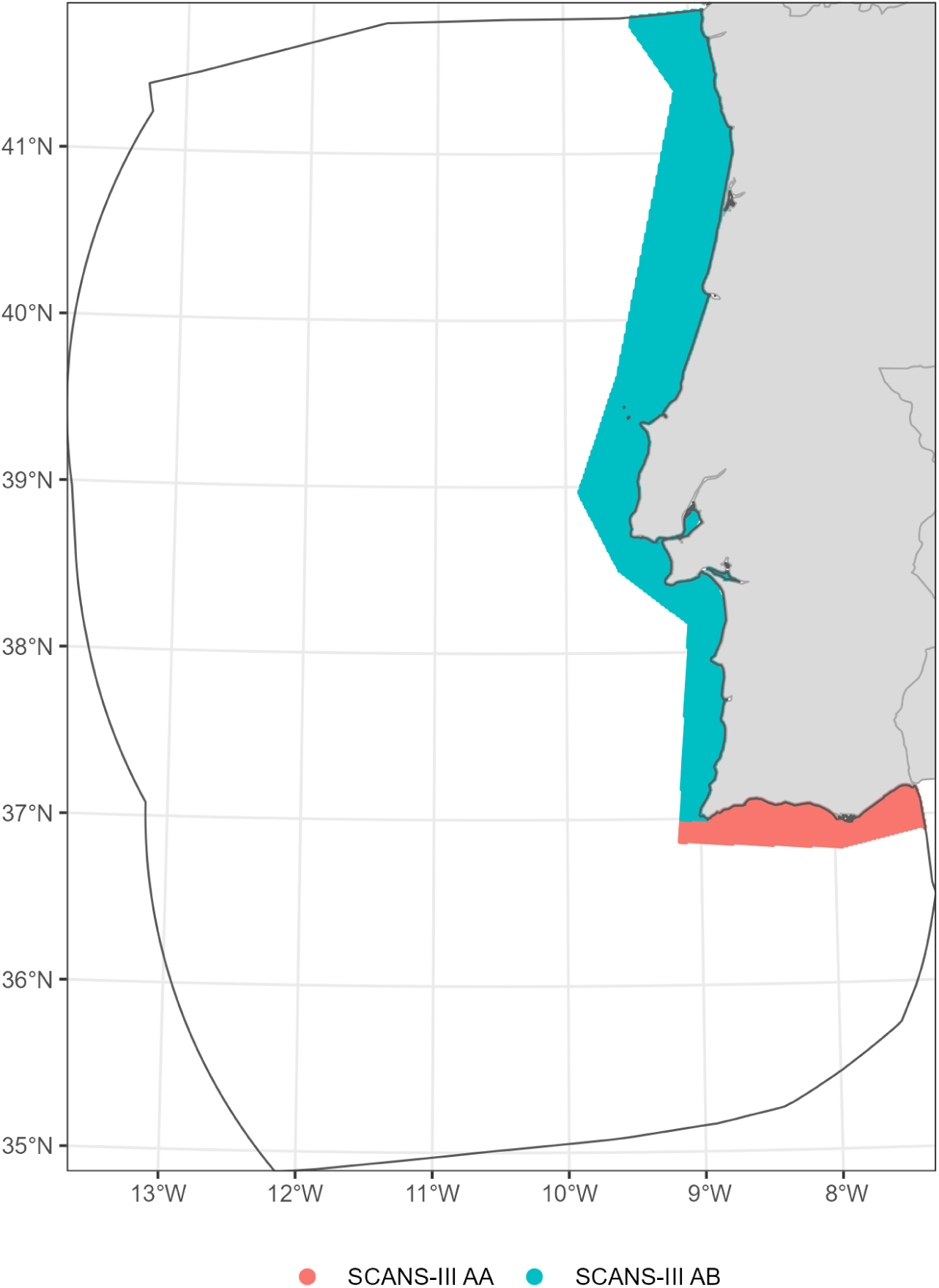
The SCANS-III survey blocks AA and AB and the prediction cells entering the density comparisons in Table S10. Both blocks extend beyond the study area; densities are compared over the shaded overlap only.

**Figure S10.**
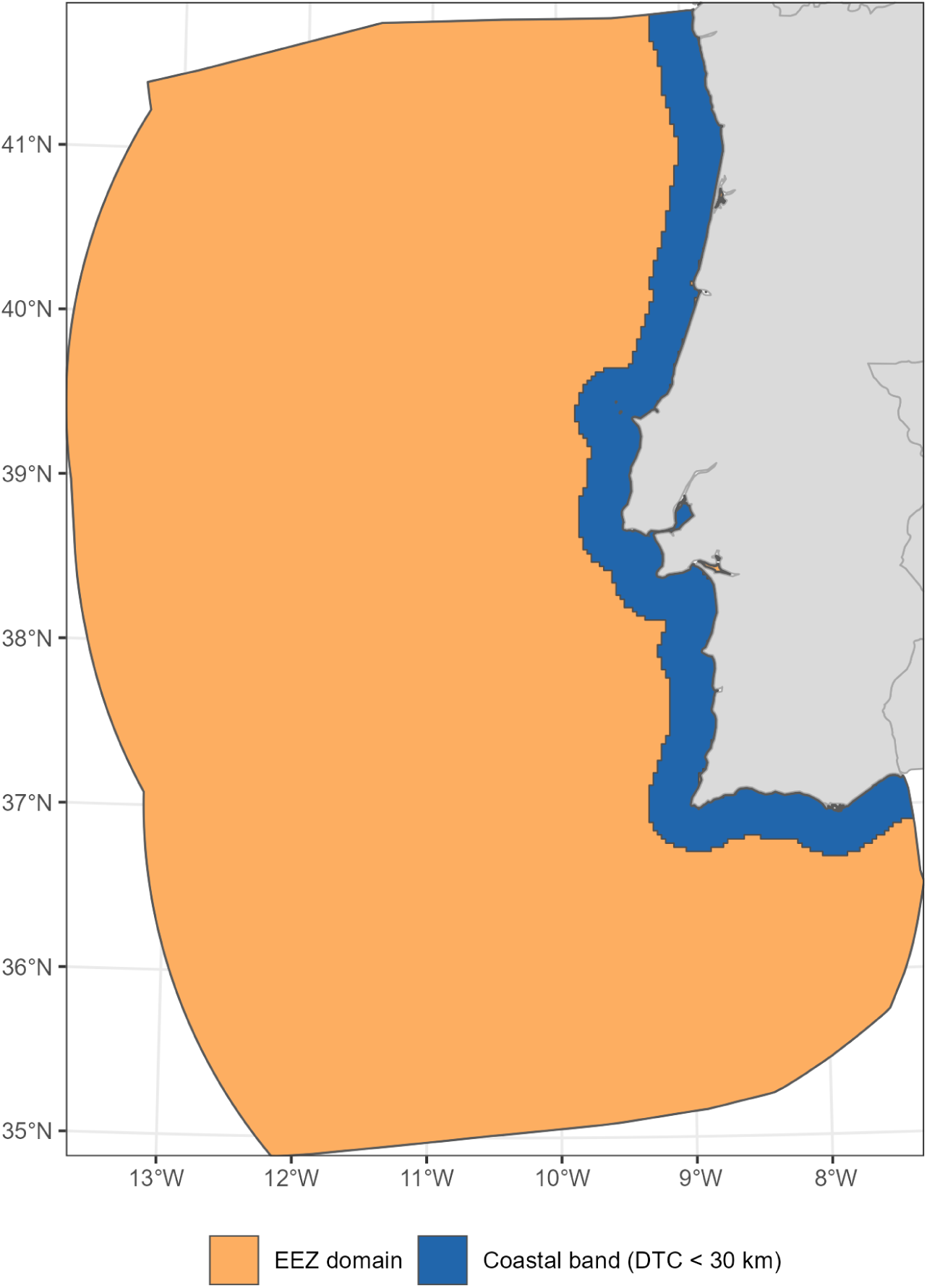
The reporting domains of Martins et al. (2026) used for the comparisons in Table S10: their EEZ domain, which corresponds to the full study area (summer and autumn; orange), and the coastal band, distance to coast < 30 km (all seasons; blue).

## Appendix S3. Mean and variance of the number of animals

Derivation supporting equation 13 of the main text.

Within a season, condition on the latent fields and hyperparameters *θ* = (*μ*(·), *κ*, . . .). Groups follow a Poisson process of intensity *λ_G_*(*s*) on *D* with Λ*_G_*=∫*_D_λ_G_*(*s*)*ds*; their sizes are themarks *G_i_*, and the number of animals is 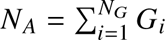.

### Mark model

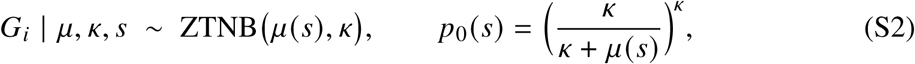

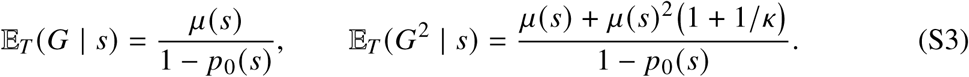

*Point process. N_G_* | *μ*, *κ* ∼ Poisson(Λ*_G_*), with locations *S* ∼ *λ_G_* (·)/Λ*_G_*.

### Mean (Campbell)

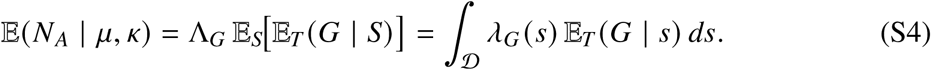

*Variance* — law of total variance over *N_G_*, marks i.i.d.:

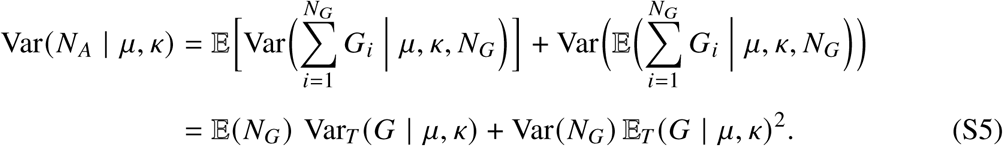

*Marginalise the mark over S* ∼ *λ_G_*/Λ*_G_:*

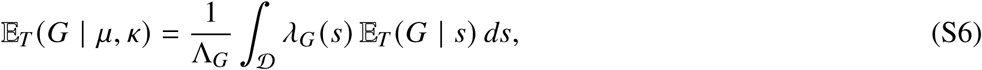

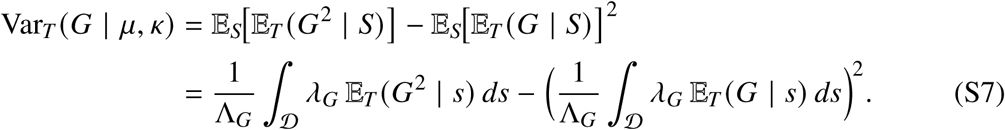

*Poisson:* E(*N_G_*) = Var(*N_G_*) = Λ*_G_*. Substituting into (S5),

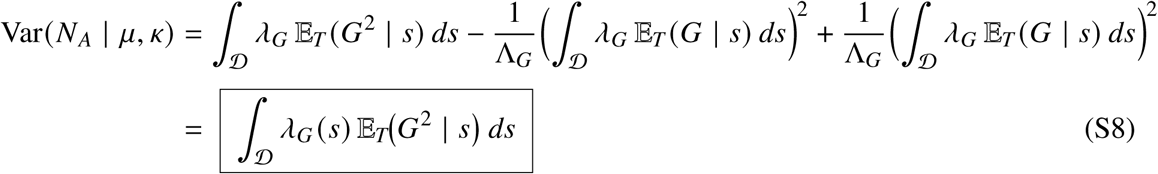

*Propagate posterior uncertainty in θ:*

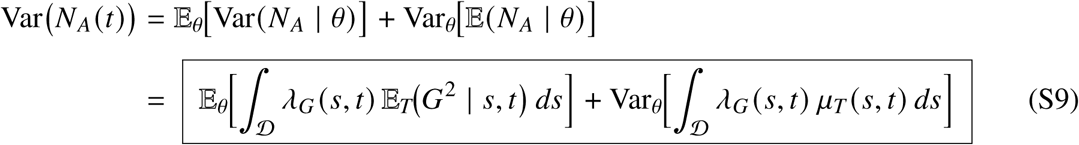

